# Aberrant activation of the innate immune sensor PKR by self dsRNA is prevented by direct interaction with ADAR1

**DOI:** 10.1101/2023.08.29.555105

**Authors:** Ketty Sinigaglia, Anna Cherian, Dragana Vukic, Janka Melicherova, Pavla Linhartova, Qiupei Du, Lisa Zerad, Stanislav Stejskal, Radek Malik, Jan Prochazka, Nadège Bondurand, Radislav Sedlacek, Mary A. O’Connell, Liam P. Keegan

**Affiliations:** Central European Institute for Technology at Masaryk University (CEITEC MU), Building E35, Kamenice 735/5, Brno, CZ 62500, Czechia; Laboratory of Embryology and Genetics of Human Malformation, Imagine Institute, INSERM UMR 1163, Université de Paris Cité, F-75015, Paris, France; Laboratory of Epigenetic Regulation, Institute of Molecular Genetics of the Czech Academy of Sciences, Vídeňská 1083, CZ 142 20, Praha 4, Czechia; Institute of Molecular Genetics of the Czech Academy of Sciences, Prumyslova 595, 252 50 Vestec, Czechia

**Keywords:** dsRNA, ADAR, RNA editing, innate immunity, PKR

## Abstract

Loss of dsRNA editing by Adar1 leads to aberrant interferon induction in *Adar* null mouse embryos. *Adar Mavs* mutants, in which this interferon induction is prevented, die within two weeks of birth. We show here that early death of pups is reduced in *Adar Mavs EIF2αk2* (Pkr) mutants, identifying PKR as the second aberrantly activated dsRNA sensor in *Adar* mutant mice. In intestines of *Adar Mavs* pups transit amplifying progenitor cells in intestinal crypts die and intestinal villi are lost in pups before death; intestinal defects are prevented in *Adar Mavs Eifak2*. In human A549 lung cancer cells ADAR1 forms a complex with PKR. AlphaFold modelling predicts a direct inhibitory interaction of ADAR1 dsRBDIII with the PKR near the kinase active site and a new mode for ADAR1 dsRNA-binding. Mutations at residues required for ADAR1 dsRNA binding or for predicted PKR contacts, prevent interaction with PKR.

---

The ADAR1 (adenosine deaminase acting on RNA 1) dsRNA editing enzyme prevents aberrant innate immune responses to cellular dsRNA. ADAR1 deaminates adenosine to inosine in dsRNA to prevent unedited endogenous cellular dsRNA from being recognized as viral. In *Adar* mutant mice the Mda5 (melanoma differentiation-associated protein 5) ^1,2^ cytoplasmic antiviral dsRNA sensor, a member of the RIG-I (retinoic acid-inducible gene I)-like dsRNA receptor (RLR) family of dsRNA receptors, is aberrantly activated and leads to aberrant induction of interferon (IFN) and to cell death. *Adar* null mutant mouse embryos lacking all Adar1 protein show aberrant high level IFN expression leading to loss of hematopoietic cells and hepatocytes in the fetal liver and die by embryonic day E12.5 ^3^. *Adar Mavs* double mutant mouse fetuses also lacking mitochondria-associated antiviral signaling (Mavs) adaptor protein required downstream of activated Mda5, avoid aberrant IFN induction and survive till birth ^1^.

Loss of inosine in dsRNA is the major factor driving aberrant Mda5 activation in the *Adar* null mutant; however, some effect of removing Adar1 protein itself also contributes. Embryos homozygous for the *Adar^E^*^912^*^A^* allele encoding an Adar1 E912A catalytically inactive adenosine deaminase that binds dsRNA, also die as embryos but survive till up to E14.5 ^2^, two days later than *Adar* null mutants. *Adar^E^*^912^*^A^ Ifih1* (Mda5) double mutant pups lacking Mda5 are viable and have normal lifespans ^2^. This was surprising because *Adar Ifih1* double null mutants die very soon after birth and it suggests that Adar1 protein also has some editing independent function ^2,4^.

The partially rescued *Adar Mavs* double null mutant pups are small and die within two weeks of birth^1^, even though aberrant IFN expression is prevented. One possible explanation for the incomplete rescue is that *Adar Mavs* pups still have another defect that may be related to the loss of Adar1p150, such as an aberrant activation of another innate immune dsRNA sensor. Another explanation, not mutually exclusive with the first one, is that *Adar Mavs* is similar to the *Adar^p^*^110^–specific knockout and has some residual, perhaps developmental, defects due to loss of Adar1p110.

Studies on cell death following *ADAR* knockout in cancer cells show that, in addition to aberrant activation of MDA5, there is aberrant activation of the dsRNA-activated protein kinase R (PKR)^5^. In *ADAR* mutant cancer cells, the aberrant PKR activation is prevented by expressing catalytically inactive ADAR1 E912A, which reduces phosphorylation of PKR. Such editing-independent effects of ADAR1 could be due to ADAR1 interactions with PKR. However, only some cancer cell lines are ADAR1 dependent ^5^. Therefore, while the role of PKR in rescuing ADAR1 deficiency has been established in some cell lines, its role in relation to ADAR1 in a whole organism is unclear. Neither MDA5 nor MAVS knockouts alone can rescue the cell viability of *ADAR* mutant cancer cells, even though second mutations removing Mavs or Mda5 prevent lethality of *Adar* mutant mouse embryos. It is imperative to fully elucidate the biological pathways that ADAR1 participates in, in a model organism; this information is essential for the development of ADAR1 as a therapeutic target for cancer treatments.

Mouse *Adar^p^*^150^–specific knockouts also have aberrant IFN induction preventable by second mutations removing Mda5 or Mavs ^6^. ADAR1p150 is also expressed constitutively in thymus and spleen. ADAR1 has an IFN-inducible p150 isoform which can shuttle into the nucleus and accumulates in the cytoplasm whereas the constitutive p110 isoform shuttles but predominantly localizes in the nucleus ^7^. Mammalian ADAR1 p150 comprises of two N-terminal Z DNA/RNA-binding domains (ZBDs) of which Zα has a nuclear export sequence ^8^, three dsRNA binding domains (dsRBDs), of which dsRBDIII has a nuclear localization sequence ^9^ and a large C-terminal adenosine deaminase domain. The first ZBD, Zα, is present only in ADAR1p150. On the other hand, *Adar^p^*^110^–specific knockout mice survive to birth, eighty percent of pups die soon after birth and twenty percent survive long-term ^10^; Adar1p110 has been proposed to play a more developmental role than Adar1 p150 ^6^.

Removing Pkr rescues aberrant IFN induction and aberrant integrated stress response (ISR) induction in a mouse *Adar^P^*^195^*^A/p^*^150^ heterozygous mutant corresponding to the homozygous *ADAR^P^*^193^*^A^* Zα domain mutation found in Aicardi-Goutières Syndrome (AGS) patients ^11^. The generation of mice with mutations in the Adar1 Zα domain are important to establish it as a model for AGS ^11–14^, however they do not reveal the consequences of a complete null mutation; one lacking the entire ADAR protein, will have on an organism. Differences between *Adar* null mutant phenotypes in those reports are likely due to different *Adar* ‘null’ mutants used, some still generate Adar1p110 or truncated Adar1 proteins ^15^ (Reviewed in ^16^). ADAR1 has RNA editing independent functions, so while a mutant may be null for RNA editing activity, it may still have an editing-independent effect that is not being measured.

Here we show that in the early-dying *Adar Mavs* double null mutant mouse pups, lacking Adar1 protein, severe intestinal defects arise by fourteen days after birth (P14). Intestinal villi are lost due to TUNEL-positive cell death leading to almost complete loss of transit amplifying stem cells (TACs), proliferating cells in the intestinal crypts that are the progenitors of the rapidly turned over differentiated absorptive and secretory cells forming the intestinal villi; quiescent stem cells do not die and are increased in numbers. In *Adar Mavs EIF2αk2* triple mutant mice also lacking Pkr, sixty percent of pups escape early death and cell death in intestinal crypts at P14 is prevented. We used Alphafold Multimer to predict that ADAR1 dsRBDIII interacts with the PKR kinase domain. We mutated predicted PKR-contacting residues in ADAR1 dsRBDIII and showed loss of ADAR1-PKR interaction. We conclude that ADAR1 interacts directly with the innate immune sensor PKR to prevent aberrant activation of PKR.

## Results

### *Adar Mavs* double mutant mouse pup short lifespan and loss of intestinal villi are rescued in *Adar, Mavs, EIF2αk2* mice lacking Pkr

The *Adar^Δ^*^2–13^ deletion mutant used here (referred to hereafter as *Adar*), removes exons two to thirteen inclusive, and deletes almost all coding sequences for both Adar1 p110 and p150, including the Adar1 p150 N-terminal Zα domain ^17^. To understand which *Adar* null mutant defects are due to loss of Adar1 protein itself, rather than to loss of RNA editing activity, we investigated residual *Adar* mutant defects still present in *Adar Mavs* double mutant pups that are born because aberrant IFN induction in embryos is prevented. *Adar Mavs* mutant pups rescued to live birth have reduced size (Fig. 1A) and Kaplan-Meier survival plots show that they mostly die within two weeks of birth, unlike the single *Mavs* mutant or wild-type C57Bl6N mice (Fig. 1B). *Adar Mavs* homozygous mutant pups weigh significantly less than their *Adar ^+/+^ Mavs* and *Adar^+/-^ Mavs* siblings and, unlike their siblings, the *Adar Mavs* homozygotes have little weight gain between P8 and P14 (Fig. 1C). To establish why *Adar Mavs* pups are small and fail to grow, we examined their internal organs (Table 1), and observed significant defects primarily in their intestines (Fig. 1E), which are shorter and narrower than wildtype at postnatal day P14 (Fig. S1A). *Adar Mavs* intestines show shortened and disrupted villi in haematoxylin and eosin-stained longitudinal sections of postnatal day 14 (P14) (Fig. 1E), proximal intestine. Intestinal villi defects have previously been reported in *Adar^p1^*^50^ mutant mice lacking the IFN-inducible isoform ^6^. Reduced white cell masses, which stain red with eosin, are also observed in spleens (Fig. S1B), consistent with previous reports of reduced white blood cells in *Adar* mutant mice ^6^.

**Figure 1.**
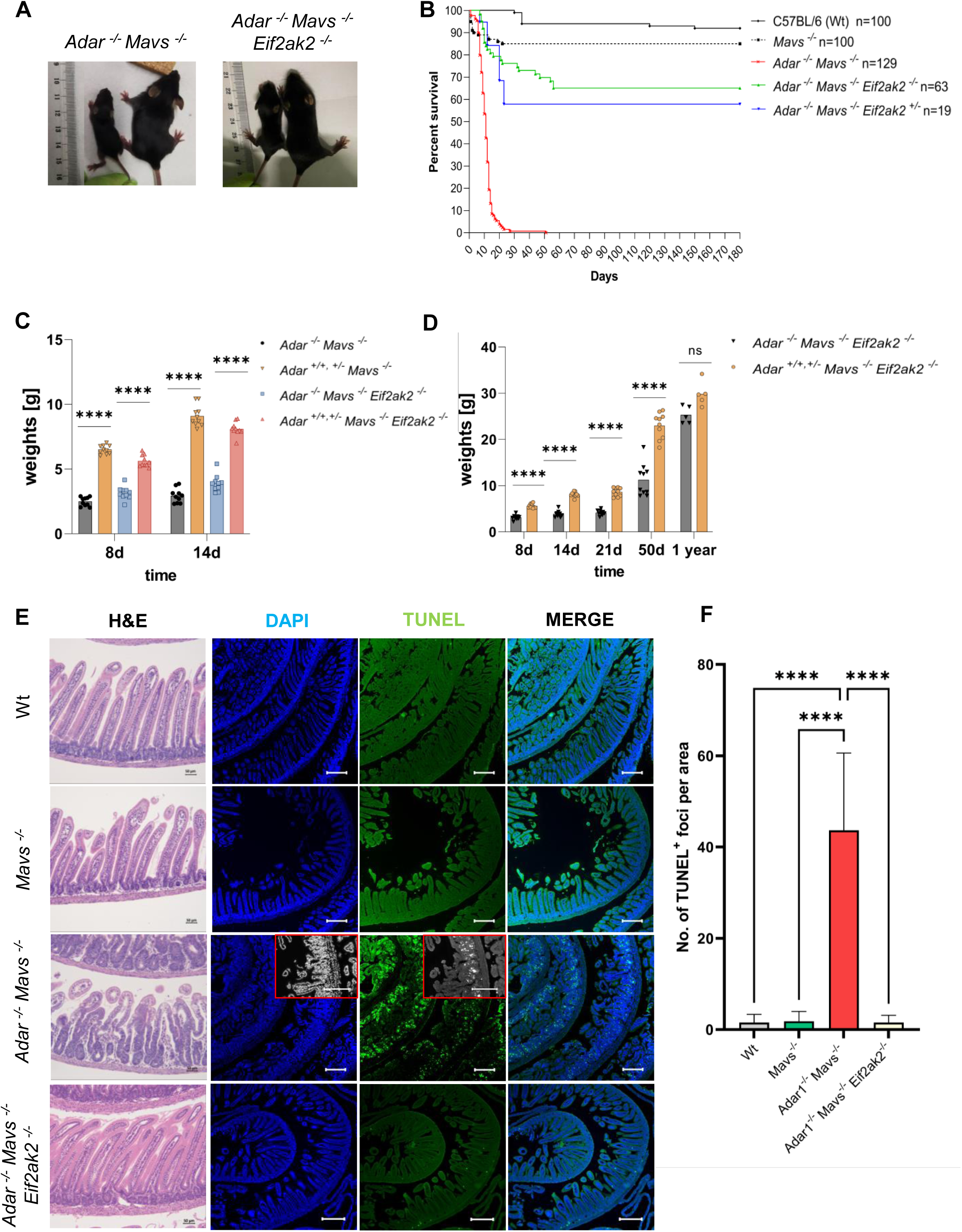
Rescue of *Adar Mavs* null mutant defects in *Adar1, Mavs, EIF2αk2* mutants lacking PKR. A. Left panel. *Adar Mavs* mutant pups (left) are small compared to their *Mavs* single mutant siblings (right). Right panel. Size is substantially rescued in *Adar Mavs EIF2αk2* mutant pups (left) versus their *Mavs EIF2αk2* siblings (right). All pups shown are 14 days old. **B.** Kaplan-Meyer plot showing that *Adar Mavs* pups die early compared to *Mavs* mutant siblings or C57Bl6 wildtype (WT). Approximately 65% of *Adar Mavs EIF2αk2* homozygous mutant pups, or 55% *Adar Mavs EIF2αk2*^-/+^ survive long-term. **C.** *Adar Mavs* mutant pups weigh less than their *Mavs* mutant siblings and do not increase in weight much between eight and fourteen days. Size is substantially rescued in *Adar Mavs EIF2αk2* mutant pups also weigh less than their *Mavs EIF2αk2* siblings but they increase in weight more than *Adar Mavs* mutant pups. **D**. *Adar Mavs EIF2αk2* mutant pups that avoid early death continue to increase in weight. **E.** Disruption of intestinal villi and cell death occurs in intestines of *Adar Mavs* mutant pups but not in *Mavs* mutant pups. Hematoxylin– and eosin-stained paraffin sections show disruption of intestinal villar structure in proximal intestine (ileum) of *Adar Mavs* mutant compared to *Mavs* mutant or C57Bl6 wildtype (WT) at P14. Proximal intestine structure is normal in *Adar Mavs EIF2αk2* mutant pups. The remaining columns show that TUNEL-positive cell death occurs in proximal intestine of *Adar Mavs* mutant pups at postnatal day P14 and is prevented in *Adar Mavs EIF2αk2* mutant pups. Scale bar is 50 μm. Images show TUNEL-positive cells (in green), DAPI staining (in blue) of nuclei and merge. Enlargements show TUNEL-positive cells more clearly in grayscale images of DAPI and TUNEL staining in *Adar Mavs* intestines. **F.** Quantitation of TUNEL-positive cells in sections of P14 intestines of *Adar Mavs* and other genotypes. TUNNEL staining was performed on three biological replicas. The statistical analysis, consisting of a one-way ANOVA followed by Kruskal-Wallis one-way analysis of variance, was performed using GraphPad Prism (version 9) and statistical significance is marked as ****p < 0.0001. Scale bar is 200 μm.

**Table 1.** Summary of the *Adar Mavs* and *Adar Mavs EIF2αk2* P14 mutant pup phenotypes.

|  | *Adar*<sup>-/-</sup> *Mavs*<sup>-/-</sup> | *Adar*<sup>-/-</sup> *Mavs*<sup>-/-</sup> *EIF2ak2*<sup>-/-</sup> |
| --- | --- | --- |
| Small Intestine | Diffuse villus atrophy;<br>Crypt, mucosa hyperplasia and vacuolation.<br>Increased apoptosis;<br>Paneth and Goblet cell hyperplasia;<br>Less and mislocated TACs;<br>Stem cell number increased with crypt hyperplasia;<br>Nucleated hematopoietic cell infiltration;<br>ISG and ISR transcripts level increased. | Phenotypically rescued.<br>Some ISG and ISR transcripts still slightly elevated. |
| Colon | No abnormalities | No abnormalities |
| Spleen | White pulp hypoplasia;<br>increased cellularity of red pulp. | Not full rescued |
| Kidney | No abnormalities | No abnormalities |
| Liver | No abnormalities | No abnormalities |
| Heart | No abnormalities | No abnormalities |

Hematoxlin– and eosin-stained 2.5 μM-thick paraffin-embedded sections through ‘Swiss rolls” ^18^, of postnatal day P14 *Adar Mavs* mutant intestines show that severe shortening of intestinal villi (Fig. 1E) occurs across proximal, medial and distal intestines at this stage. The intestine comprises of duodenum, jejunum and ileum in anterior to posterior order. However, these anatomical subdivisions are of very unequal lengths and are difficult to distinguish in intestines of pups, so we have divided pup intestines in three equal proximal, medial and distal sections and processed these in parallel in these experiments. TUNEL staining to detect dying cells in 2.5 μM-thick paraffin sections of *Adar Mavs* double mutant intestines show major death of cells in P14 intestinal crypts (Fig. 1E,F), where intestinal stem cells divide to generate cellular precursors for renewal of intestinal villi.

Intestinal defects are progressive in pups after birth; no intestinal cell death was observed in *Adar Mavs* fetuses at E16.5 (Fig. S1C) nor in intestines of pups at birth (P0) (Fig. S1,D). Intestines from postnatal day P8 pups showed defects in villi and TUNEL positive cell death only in proximal and medial intestine (Fig. 2A,B) and not in distal intestines. This observation is in agreement with a previous report ^6^ but very dissimilar to another describing TUNEL positive cells in the ilea of an *Adar Mavs* ^12^ line different from ours; the difference in the latter case can likely be attributed to that *Adar* mutant still expressing Adar1 p110 isoform^15^. Thus, we conclude that in a complete null for Adar1, TUNEL-positive cell death in *Adar Mavs* intestines begins in proximal intestine by postnatal day P8 and progresses to the whole intestine by P14. This progression of intestinal defects occurs over the days leading up to deaths of *Adar Mavs* mice as indicated in the Kaplan Meier plot (Fig. 1B).

**Figure 2.**
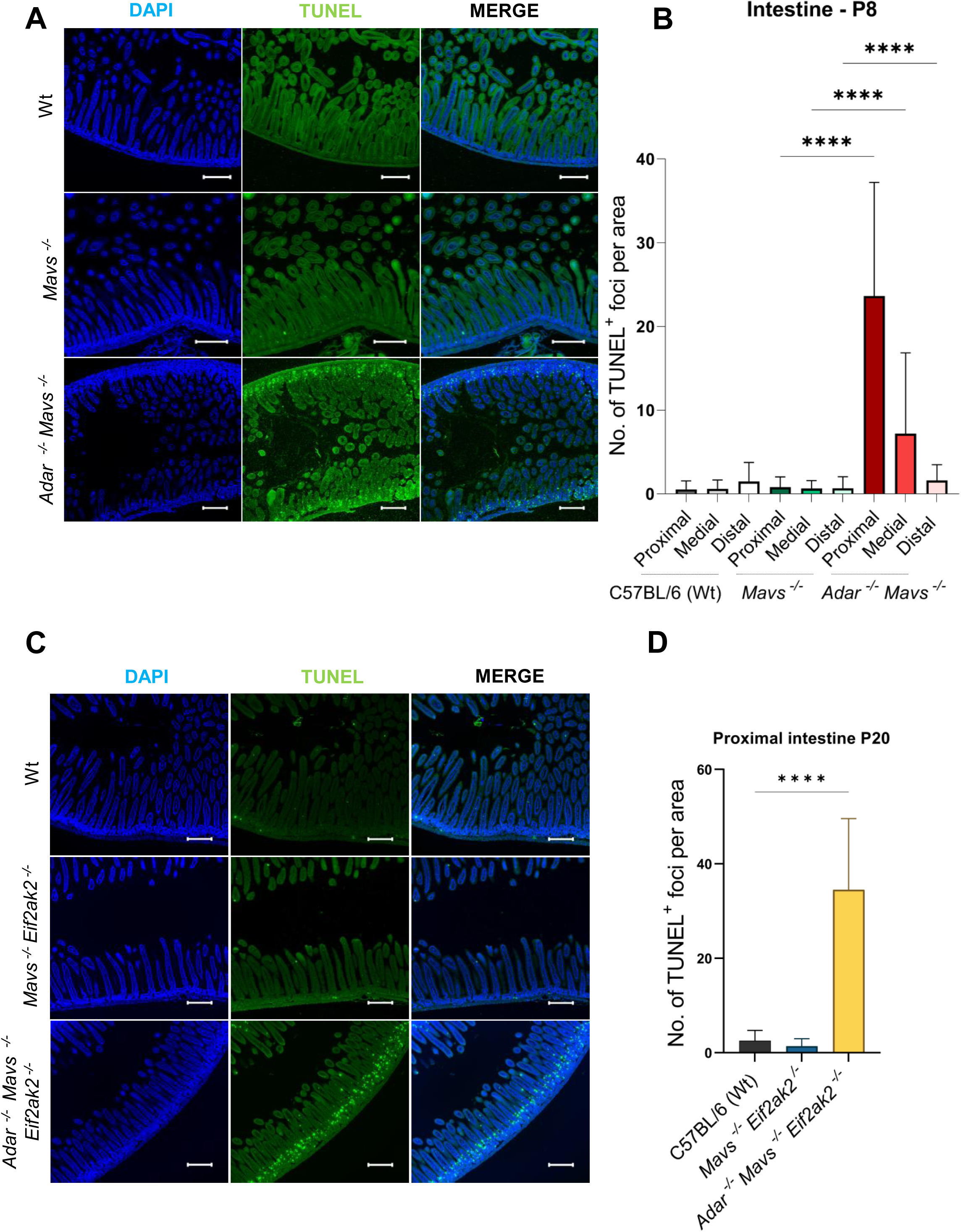
TUNEL-positive death of intestinal crypt cells in *Adar Mavs* proximal intestine begins by P8, extends throughout small intestines and colon by P14 and is postponed in *Adar Mavs EIF2αk2* triple mutants. A. TUNEL-positive cell death in *Adar Mavs* double mutant P8 proximal intestines compared to *Mavs* mutant or C57Bl6 wildtype (WT). The 200 μm scale bar is indicated in white. **B.** Quantitation of TUNEL-positive cells in proximal, medial (jejunum) and distal intestine (duodenum) of C57Bl6 wildtype (WT), Adar Mavs double mutant and Mavs at P8. **C**. TUNEL-positive cell death in *Adar Mavs* double mutant P20 proximal intestines compared to C57Bl6 wildtype (WT) or *Adar Mavs EIF2αk2* triple mutants; note that TUNEL-positive cell death reappears at 20 days in *Adar Mavs EIF2αk2* in cells in the villi rather than in the crypts. The 200 μm scale bar is indicated in white. **D.** Quantitation of TUNEL-positive cells in P20 proximal intestine of C57Bl6 wildtype (WT), *Adar Mavs EIF2αk2* triple mutant and their *Adar Mavs* double mutant siblings. TUNNEL staining was performed on three biological replicas. The statistical analysis, consisting of a one-way ANOVA followed by Kruskal-Wallis one-way analysis of variance, was performed using GraphPad Prism (version 9) and statistical significance is marked as ****p < 0.0001.

To determine whether aberrant activation of the dsRNA sensor Pkr contributes to the death of *Adar Mavs* pups, we used a new *EIF2αk2* (Pkr) null mutant line to generate *Adar Mavs EIF2αk2* triple mutants. The small size of *Adar Mavs* mutant pups is partially rescued in *Adar Mavs EIF2αk2* triple mutant pups at birth and, unlike *Adar Mavs* pups, they continue growing slowly (Fig. 1C) and approach normal sizes (Fig. 1A) and weights (Fig. 1D) over time. The Kaplan-Meier plot of survival of *Adar1 Mavs EIF2αk2* mutants shows that *Adar Mavs* mutant early death is also substantially prevented, with sixt-five percent of *Adar1 Mavs EIF2αk2* mutant animals surviving beyond one month and showing long-term survival thereafter (Fig. 1b). In fact, even *Adar Mavs EIF2αk2^-/+^* heterozygotes with half the normal dosage of Pkr are rescued to sixty percent, indicating strong Pkr dosage sensitivity (Fig. 1B). Once the pups are past weaning, no further death is observed. We observed no difference between the spleen of *Adar Mavs* pups and *Adar Mavs EIF2αk2* triple mutants (Fig. S1B), so there is no evidence that the spleen phenotype is relevant to the observed early lethality of *Adar Mavs* mutants. Instead, we focused on the intestines of *Adar Mavs, EIF2α2* mutants and observed that gut lengths and thickness look more like wildtype (Fig. S1A). Haematoxylin– and eosin-stained paraffin sections displayed restoration of normal intestinal villi in *Adar1 Mavs EIF2αk2* mutants (Fig. 1E,F), and prevention of TUNEL-positive cell death in intestines (Fig. 1E,F). This is consistent with the idea that the intestinal defects arising due to aberrant Pkr activation in *Adar Mavs* cause the deaths of these mice.

Surprisingly, we find that, even though the intestinal villi are closer to normal in *Adar1 Mavs EIF2αk2* than in *Adar1 Mavs,* nevertheless TUNEL-positive cell death reappears in *Adar1 Mavs EIF2αk2* when examined in older pups, at P20 (Fig. 2C, D), even though half of these mice survive and continue to grow past P20 (Fig. 1D). We note, however, that the TUNEL-positive cells are now observed at a more apical position than in *Adar Mavs* mutants, are less severe and occur lower in the villi rather than in intestinal crypts.

### Loss of Ki-67 positive transit-amplifying progenitor cells in intestinal crypts in *Adar Mavs* and aberrant differentiation of secretory cells types are rescued in *Adar, Mavs, EIF2αk2*

The severe intestinal defects in *Adar Mavs* double mutant pups might result from loss of stem cells in the intestinal crypts. However, quiescent stem cells in the bases of the crypts appeared similar to these stem cells in wildtype or *Mavs* mutant intestines, as detected by staining for the stem cell marker Olfm4 (Fig. 3A,B). However, the number of Olfm-positive quiescent stem cells is increased in *Adar Mavs* intestines (Fig. 3A,B).

**Figure 3.**
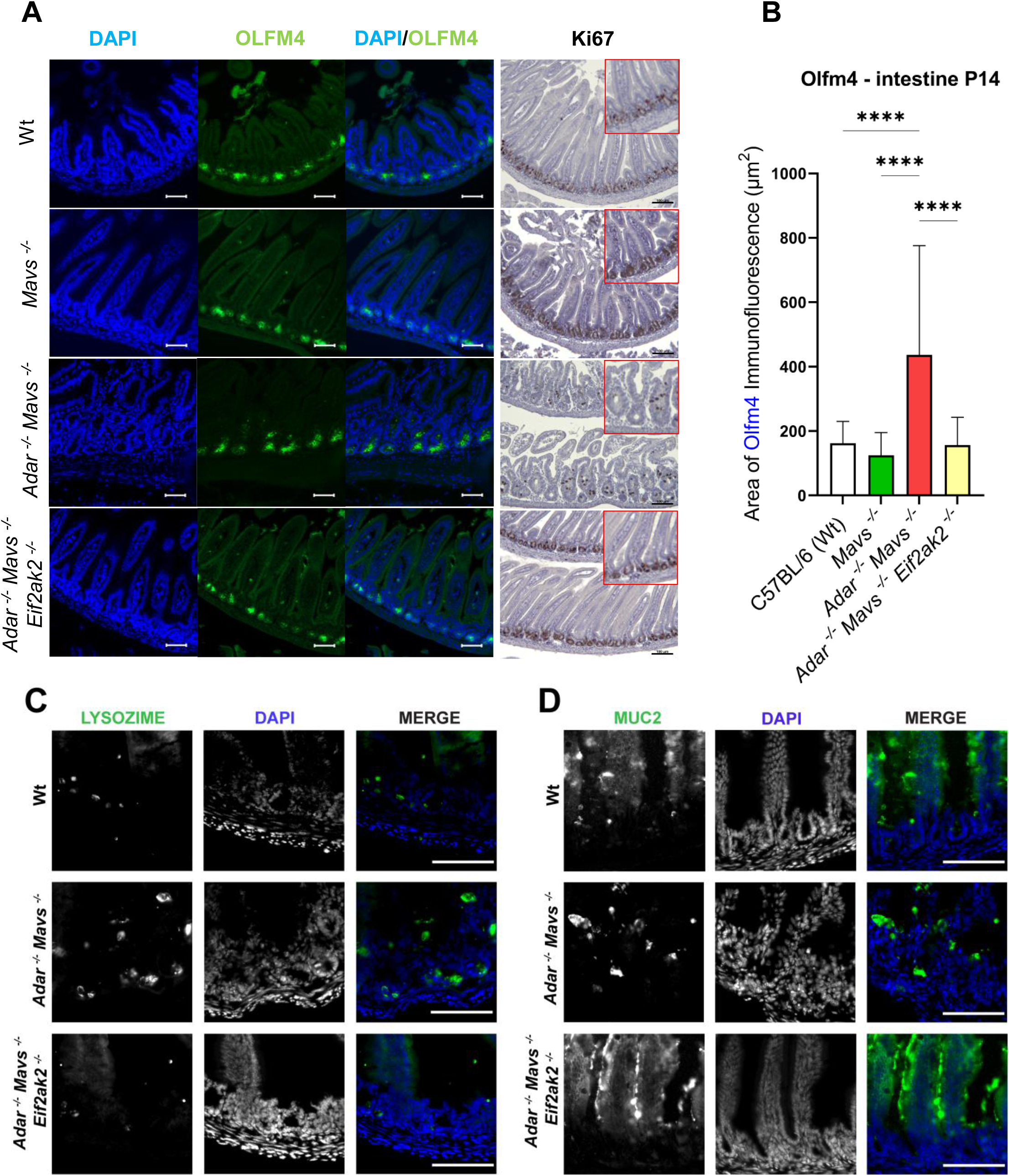
*Adar Mavs* double mutant intestines show almost total death of transit amplifying progenitor cell (TACs); cell death is delayed and occurs in villi in *Adar Mavs EIF2αk2* triple mutants. A. Left columns. Olfm4-positive intestinal stem cells in the bases of the intestinal crypts are increased in *Adar Mavs* P14 double mutant intestines compared to C57Bl6 wildtype (Wt) or *Mavs* mutant intestines and are normal in *Adar Mavs EIF2αk2* triple mutants. Columns show DAPI, Olfm4 and merge. The 50 μm scale bar is indicated in white. Right column. Loss of Ki-67-positive transit amplifying cells (TACs) occurs in *Adar Mavs* intestines and is rescued in the *Adar Mavs EIF2αk2* triple mutant. The 100 μm scale bar is indicated in white**. B.** Quantitation of Olfm4 staining in wt, *Mavs*, *Adar Mavs* and *Adar Mavs EIF2αk2* proximal intestines at P14. **C.** Paneth Cells, which are normally located near the intestinal stem cells in the bases of the intestinal crypts in Wt and in *Adar Mavs EIF2αk2* intestines are aberrantly localized more apically in *Adar Mavs*, in large cells in the villi. Columns of images show anti-Lysozyme staining, DAPI staining and merge. The 100 μm scale bar is indicated in white. **D**. Mucin two (MUC2), a heavily glycosylated protein secreted from Goblet cells to form mucus, is expressed in a punctate pattern in the *Adar Mavs* P14 proximal intestine whereas it is spread normally along the surfaces of the villi in wt and *Adar Mavs EIF2αk2* intestines. Staining in panels A and B were performed on three biological replicas, two biological replicas were used in panels C and D. The 100 μm scale bar is indicated in white.

Since quiescent stem cells were not decreased in *Adar Mavs* intestines, we next examined the immediate progeny of the quiescent stem cells, the rapidly proliferating transit amplifying cells (TACs) immediately above the quiescent stem cells in the intestinal crypts. We used immunostaining against Ki-67 in cryosections of *Adar Mavs* intestines to examine the status of TAC progenitor cells (Fig. 3A). The Ki-67 staining in TAC cells in wildtype and *Mavs* mutant intestines (Fig. 3A), reflects expected cell cycle changes in expression and localization, with some cells showing more dispersed Ki-67 staining while others show staining in nucleoli (Fig. 3A). Ki-67 is strongly expressed in dividing stem cells ^19^, and is strongly downregulated in nondividing G0 cells; during G1 and G2 the Ki-67 protein localizes to nucleoli whereas, during mitosis, Ki-67 covers chromosome arms ^20^. Much fewer Ki-67-positive TAC cells are observed in *Adar Mavs* P14 intestines (Fig. 3A) compared to wild type or *Mavs* mutant intestines. The intestinal cell death in *Adar Mavs* is most similar to that in an Lgr5-positve intestinal crypt stem cell specific knockout of *Adar* in which quiescent stem cells and their TAC progeny all die ^21^; prevention of the aberrant interferon induction in *Adar Mavs* evidently allows these cells to survive to later stages.

In *Adar Mavs* intestines the few surviving Ki-67 positive cells are more apical than normal and larger, with Ki-67 staining always located in what appear to be enlarged nucleoli (Fig. 3a). The reduced number and altered appearances of Ki-67 positive TACs are rescued in *Adar Mavs EIF2αk2*. Surprisingly, TUNEL-positive cell death appears again in *Adar1 Mavs EIF2αk2* when examined at P20 (Fig. 2d) although it appears to occur higher up in the intestinal crypts and in the villi themselves, as though cell death is now occurring later in TAC proliferation and perhaps not eliminating them completely, allowing maintenance of villi. Evidently, a change in the pattern of cell death, which allows maintenance of some proliferating progenitor TAC cells underlies the rescue in *Adar1 Mavs EIF2αk2*.

Further indications that TAC progenitor cells are abnormal in *Adar Mavs* intestines come from examination of Paneth cells and Goblet cells, the main secretory cell progeny of the TAC progenitor cells. Paneth cells, normally differentiate from the TACs and then migrate back down among the quiescent basal stem cells in the crypts, where they secrete both antimicrobial peptides and signals for maintenance of quiescent stem cells. Staining intestinal paraffin sections of wildtype or *Mavs* intestines with anti-Lysozyme antibodies detects normal Paneth cells in intestinal crypts (Fig. 3C). However, in *Adar Mavs* intestines Lysozyme-positive cells show abnormally increased size and aberrantly apical locations (Fig. 3C). This *Adar Mavs* Paneth cell defect is corrected in *Adar Mavs EIF2αk2* triple mutant intestines (Fig. 3C). Clearly, differentiation of TAC progenitor cells is abnormal and failure to produce normal Paneth cells that migrate back down into the crypts may contribute to the abnormal increase in the number of quiescent stem cells in the crypts through a feedback effect.

The other type of secretory cell, Goblet cells in the villi, produce mucin that hydrates and protects villi. Anti-Mucin2 (Fig.3D) staining of intestinal sections shows normal secreted Mucin2 along the sides of the villi in wildtype or *Mavs*; however, *Adar Mavs* villi show reduced secretion of the Mucin2 glycoprotein (Fig. 3D), with possible retention of Mucin2 in large secretory vesicles in aberrantly enlarged cells. These enlarged cells could be same aberrant secretory cells detected by anti-Lysozyme staining in *Adar Mavs* (Fig. 3C) or even the same cells as the aberrant Ki-67 positive cell (Fig. 3A). Aberrant anti-Muc2 staining in *Adar Mavs* is also rescued in *Adar Mavs EIF2αk2* triple mutants (Fig. 3D). In summary, abnormal differentiation of Ki-67 positive TACs in the *Adar Mavs* crypts is fully rescued in *Adar Mavs EIF2αk2* (Pkr) triple mutants (Fig. 3 A).

We examined gut microbiomes in pups of the different genotypes but results show high variability and only one type of bacterium, which is also increased in inflammatory bowel disease, is increased in *Adar Mavs* intestines compared to wildtype (Figure S3).

### Defects in cells supporting the intestinal stem cell niche and increased local IFN and integrated stress response transcripts in *Adar Mavs* intestines are rescued in *Adar, Mavs, EIF2αk2* mutants

There is clearly a strong cell autonomous requirement for Adar1 in intestinal stem cells^21^. In *Adar Mavs* however the loss of TACs might be partly due to loss of Adar1 in surrounding cells that help maintain the intestinal stem cell niche. In the intestinal crypts, the long-term stem cells are supported and proliferation of stem cells is regulated by, among other signaling molecules, Wnt proteins secreted from enteric glial cells (EGCs) which surround the base of the crypt ^22^. EGCs, like enteric neurons, originate from embryonic neural crest cells that migrate into the developing gut between E10.5 and E14. Long considered bystanders providing support to neurons, EGCs have emerged as a signaling hub capable of responding to environmental and tissue changes by producing diffusible factors that modulate the behavior of surrounding cells ^22^. In appearance EGCs resemble astrocytes but they also resemble innate immune cells because they secrete immune cytokines in response to microbiome stimulation, pathogen invasion and inflammation or tissue injury^22^.

In the *Adar Mavs* double mutant embryos, neural crest derivatives such as EGCs, might have some residual defect arising from the loss of Adar1 protein that has not been fully prevented in the *Adar Mavs* double mutant. To determine whether EGCs or enteric neurons are abberant, we stained cryosections of *Adar Mavs* double mutant intestines with antibodies against Glial Fibrillary Acidic Protein (GFAP) (Fig. 4A) or TUJ1 neuron-specific class III beta-tubulin (Fig.S2A) to detect activated EGCs or neurons, respectively. TUJ1-positive neurons appear normal (Fig. S2A) as do anti-SMA stains of muscle layers (Fig.S2B). EGCs expressing GFAP, which is also induced by inflammation (GFAP+ EGCs), are usually located mainly in two plexus layers, the sub-mucosal plexus layer beneath the intestinal crypts in the mucosa and the much deeper myenteric plexus layer located between the circular and longitudinal muscle layers. In *Adar Mavs* intestines the GFAP+ staining appears reduced in the submucosal plexus layer and instead extends up higher around the crypts and villi (Fig. 4A); CD45-positive cells are also recruited here (Fig. 4B), consistent with ongoing inflammation in this area. The aberrant GFAP staining in villi is absent, and CD45 staining is similar to controls in *Adar Mavs EIF2αk2* triple mutant intestines (Fig. 4 A,B), suggesting that inflammatory effects are reduced to a least wild-type levels. Staining with antibodies against epithelial E-cadherin also showed loss of basal E-cadherin staining in the intestinal crypts (Fig. S2C).

**Figure 4.**
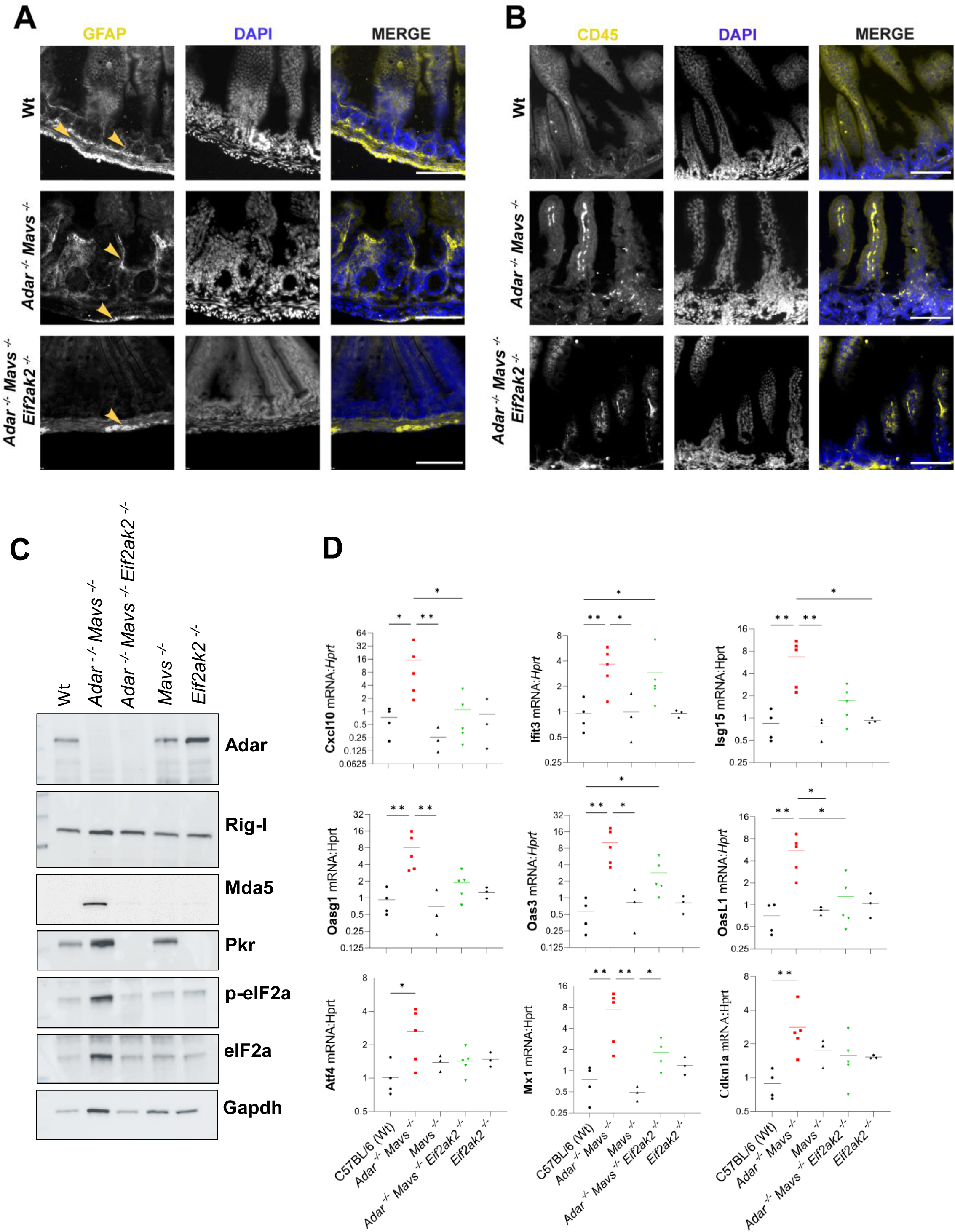
Inflammation/interferon induction and integrated stress response (ISR) induction occur in *Adar Mavs* double mutant P14 proximal intestines compared to wt or *Mavs* mutant intestines and these inductions are prevented in *Adar Mavs EIF2αk2* triple mutants. A. Increased and abnormally more apically localized expression of Glial Fibrillary Acidic Protein (GFAP, yellow) in Enteric Glial Cells (EGCs) in *Adar Mavs* intestines compared to normal expression in myenteric and submucosal plexus layers in wild-type and in *Adar Mavs EIF2αk2* intestines indicate increased inflammation in *Adar Mavs* intestines. Columns of images show anti-GFAP staining, DAPI staining and merge. The 100 μm scale bar is indicated in white. **B.** Aberrantly increased anti-CD45 staining of T-cells in villi in *Adar Mavs* intestine compared to normal anti-CD45 T cell staining in Wt and *Adar Mavs EIF2αk2* intestines. Columns of images show anti-CD45 staining, DAPI staining and merged images. The 100 μm scale bar is indicated in white. Two biological replicas were used in panels A and B. **C.** Immunoblot analysis of protein extracts from P13 proximal intestines from mouse pups of different genotypes. **D.** RT-PCR analyses of interferon-induced (*Cxcl10, Isg15, Ifit3, Oas3, Oas1g* and *OasL1*) and integrated stress response (ISR) (*Atf4, Mx1* and *Cdkn1a*) transcript expression in proximal intestines of P14 pups of C57Bl6 (Wt), *Adar Mavs*, *Mavs*, *Adar Mavs EIF2αk2* and *EIF2αk2* genotypes.

Immunoblot analyses of total protein extracts from proximal intestines of P14 pups show that *Adar Mavs* intestines have increased expression of Pkr, Mda5 and Rig-I proteins compared to wildtype or *Mavs* intestines (Fig.4C); in the absence of RLR signaling due the removal of Mavs this interferon induction might be through Toll receptors. Pkr is also somewhat elevated in *Mavs* intestine (Fig.4C); the Mavs mutant has been reported to cause some sterile inflammation in the brain ^23^. We cannot assess the activation state of Pkr protein in mice because antibodies against active, autophosphorylated pT446 PKR do not detect the mouse protein. Antibodies against the major Pkr target protein eIF2α or its phosphorylated form pS51 eIF2α, show increased expression of eIF2α in *Adar Mavs* intestines as well as increased phosphorylation. Aberrant activation of Pkr occurs in *Adar Mavs* intestine and this could cause direct activation of cell death with increased IFN or integrated stress response (ISR) induction. We examined expression of IFN stimulated gene (ISG) and ISR transcripts in *Adar Mavs* P14 proximal intestines by qPCR. There is some increased expression of ISG transcripts *Cxcl10, Oas3, Isg15, Oasg1, Mx1, Oasl1* and *Ifit3* (Fig. 4B), in *Adar Mavs* intestines compared to *Mavs* mutant or wildtype intestines and this aberrant ISG transcript expression is prevented in *Adar Mavs EIF2αk2*. However, this ISG transcript induction is relatively weak and may reflect the localized inflammation observed near dying TAC progenitor cells where increased GFAP and CD45 expression are observed. We also observe slight increases in expression of ISR transcripts *Atf4* and *Cdkn1a* specifically in the proximal intestine (Fig. 4B), in *Adar Mavs* compared to *Mavs* or wildtype, which is prevented in *Adar Mavs EIF2αk2*. Inductions of ISR transcripts in other studies are also very weak, about tenfold ^11^, and much lower than those possible for ISGs. Unlike a previous report on *Adar^P^*^195^*^A/p^*^150^ heterozygous mutant where ISR transcript induction was observed in liver and other tissues ^11^, we did not detect ISR transcript induction in those organs.

### ADAR1 interacts with PKR and ADAR1 binding to dsRNA is required for ADAR1-PKR interaction

ADAR1 has been previously reported to associate with PKR in the Jurkat T cell line where both proteins are moderately expressed, based on co-immunoprecipitation experiments. We confirmed this by immunoprecipitation on PKR from A549 lung cancer cells, which also have moderate levels of both proteins, followed by detection of ADAR1 in the immunoprecipitated sample (Fig. 5A). Both cytoplasmic ADAR1 p150 and nuclear ADAR1 p110 protein coimmunoprecipitate with PKR (Fig. 5A). PKR is mostly in the cytoplasm with up to twenty percent found in the nucleus in cell lines, so the PKR interacts with endogenous ADAR1 p150 in cells whereas PKR interaction with ADAR1 p110, which is more abundant than ADAR1 p150, may occur mainly after cell lysis when there is mixing of nuclear and cytoplasmic fractions.

**Figure 5.**
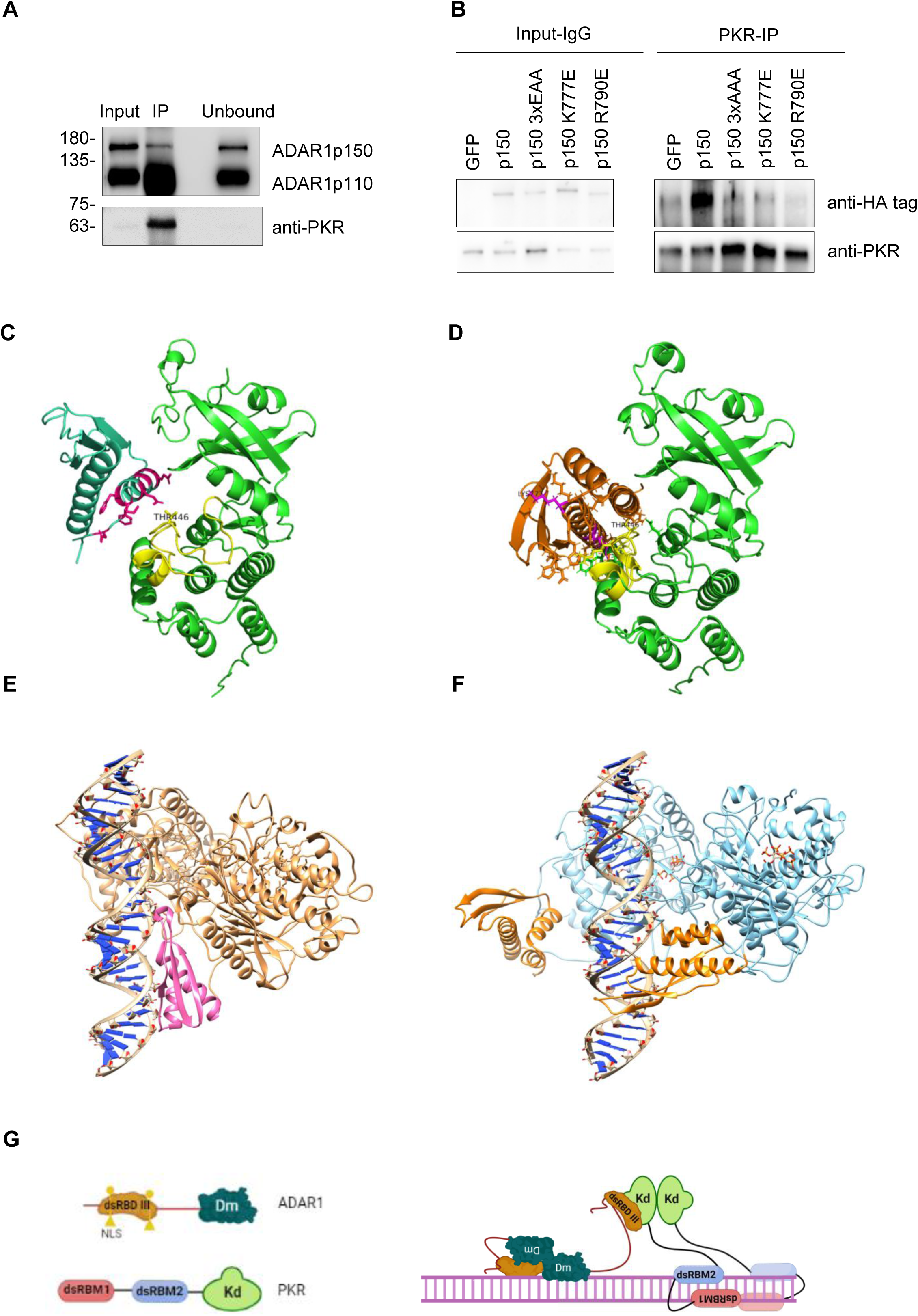
Human ADAR1 interacts directly with the kinase domain of PKR. A. Co-immunoprecipitation with PKR from A546 lung cancer cells of ADAR1 p150 and ADAR1 p110 isoforms (n=2). **B.** Co-immunoprecipitation with PKR from A546 lung cancer cells of ADAR1 p150-HA and mutant proteins expressed from transfected plasmids. The ADAR1 mutants tested are the ADAR1 dsRBDIII 3X(EAA) dsRNA-binding mutant in which the SKKxAK motif has been mutated to SEAx<u>A</u>A in all three dsRBDs and the ADAR1 K777E and ADAR1 R790E mutant proteins in which possible PKR-interacting residues in ADAR1 dsRBDIII are mutated (n=3). **C.** AlphaFold prediction of the structure of full-length human PKR showing residues at the N-terminus of helix1 of dsRBDI (in green cyan with some residue side chains indicated) interacting with the C-lobe of the kinase domain (both kinase domain lobes are in green), over the activation loop of PKR kinase domain (in yellow) with the Thr446 residue in the activation loop labelled. PKR dsRBDII and the linkers around it have been hidden for clarity. **D.** AlphaFold Multimer prediction of the ADAR1 dsRBDIII (orange) protein-protein interaction with the PKR kinase domain (green) on the C-lobe near the kinase active site at the junction between the N and C lobes. Mutated ADAR1 dsRBDIII residues interacting with PKR are in magenta. Side chains of other ADAR1 dsRBDIII residues that may contact PKR are also shown in orange. To emphasize the similarity of the PKR dsRBDI and ADAR1 dsRBDIII interactions with the PKR kinase domain these models can be viewed side by side in the attached Movie Fig. 5C,D. **E.** The ADAR2 dimer on dsRNA positions dsRBDII (in hot pink) parallel to the dsRNA for canonical dsRBD-dsRNA contacts. **F.** The predicted ADAR1 dimer on dsRNA will hold dsRBDIII (in orange) by the alpha-helical end of the dsRBD and point helix3 towards one strand of the dsRNA. The ADAR-dsRNA and ADAR1-dsRNA complexes can be viewed side by side in the attached Movie Fig. 5E,F.

To test whether the ADAR1-PKR interaction depends on ADAR1 dsRNA binding we expressed wildtype or mutant, HA-tagged versions of human cytoplasmic ADAR1 p150 in A549 lung cancer cells treated with IFN and performed immunoprecipitations of endogenous PKR protein. Immunoblot analysis of the precipitate shows that HA-ADAR1p150 co-immunoprecipitates efficiently with PKR (Fig. 5B). To test whether ADAR1-PKR interaction requires dsRNA binding to dsRNA we introduced three mutations in the S<u>KK</u>xA<u>K</u> motif at the N-terminus of the last, long, α helix in all three dsRBDs (ADAR1p150 3xEAA) ^24^. We transfected a plasmid expressing ADAR1 3xEAA into A549 cells and tested whether this ADAR1 mutant still co-immunoprecipitates with endogenous PKR The ADAR1 3xEAA mutant fails to coimmunoprecipitate with PKR, indicating that the interaction with PKR was lost (Fig. 5B). The ADAR1 3xEAA mutant is known to eliminate dsRNA-binding and ADAR1 RNA editing activity ^24^.

### Predicted interaction of ADAR1 dsRBDIII with PKR kinase domain

To examine how ADAR1 may directly interact with PKR we first examined the structures of the individual proteins in the <u>AlphaFold Protein Structure Database</u> (ASD) ^25^, of predicted full-length protein monomer structures. PKR is known to be auto-inhibited, in the absence of dsRNA, by the N-terminal dsRBDs. In the presence of dsRNA, the PKR dsRBDs bind to it and the C-terminal PKR kinase domain is then able to form a pre-activated, back-to-back (BTB) PKR dimer through the N-lobe of the kinase domain. Autophosphorylation on Thr446 in the kinase domain activation loop completes the activation of PKR. Predicted structures of PKR monomers from different vertebrates are in the ASD; human PKR shows dsRBD1 helix1 and the PKR N-terminal tail positioned on the PKR kinase domain N-lobe, right over the activation loop containing Thr446 (Fig. 5C, Movie Fig. 5C,D); rat Pkr shows a very similar positioning of helix1 of dsRBD2 instead on the kinase domain. No crystal structure is available for the full-length autoinhibited PKR monomer and the predicted PKR monomer structures probably show the autoinhibited state of PKR in the absence of dsRNA. Since PKR and ADAR1 are similar in that they have several N-terminal dsRBDs, we wondered if the autoinhibitory interaction of PKR dsRBDs with the PKR kinase domain might be succeeded by an inhibitory interaction of an ADAR1 dsRBD with the kinase domain following the same pattern.

We used a local installation of AlphaFold Multimer ^26^ to predict possible interactions between different domains of PKR and ADAR1. When interaction predictions are run for the PKR kinase domain alone with either the three ADAR1 dsRBDs together or dsRBDIII alone, AlphaFold Multimer finds a potential ADAR1 dsRBDIII interaction with the PKR kinase domain (Fig. 5D, Movie Fig. 5C,D). In these models dsRBDIII helix1 C-terminus and helix3 N-terminus associate with the PKR kinase domain C-lobe over the PKR activation loop like dsRBD1 helix1 N-terminus in the PKR monomer (Fig. 5D, Movie Fig. 5C,D). AlfaFold is unable to model interactions with dsRNA so the PKR dsRBDs must be removed manually to enable the prediction of the PKR kinase domain interaction with ADAR1 dsRBDIII. The ADAR1 monomer structure also has an interaction of dsRBDIII with the deaminase domain which must be removed manually. When ADAR1 dsRBDIII is removed, neither dsRBDI nor dsRBDII is predicted to interact with PKR. When full length ADAR1 and PKR proteins are tested for interaction only the separated monomers are predicted as in the ASD models.

AlfaFold Multimer generates a series of models for each prediction; we usually request five or ten. To examine how similar the multiple models generated for the ADAR1 dsRBDIII-PKR kinase domain interaction obtained from each AF Multimer run are to each other, we used the Chimera Matchmaker tool to superimpose the AlphaFold Multimer interaction predictions (Fig. S4A, Movie Fig. S4A). For the ADAR1 dsRBDIII – PKR kinase domain interaction we superimposed the models using the PKR kinase domain and then examined how dsRBDIII changes position in the different models. Three out of five models show very similar positions of the N-terminus of dsRBDIII helix 3 and the nearby beta sheet loop and vary mainly in the angle which helix3 makes away from this position (Fig. S4A, Movie Fig. S4A). We conclude that the predicted interaction is likely to be a true one.

After the dsRNA binding of PKR allows BTB dimerization of the kinase domain through the N-lobes, this prepares the active site region in the cleft between the N– and C-lobes of the kinase domain for the final activation step by phosphorylation of residue T442 ^27^. In PKR, phosphorylation of pT442 is required for full kinase activity because residues from the kinase domain N– and C-lobes interact with the phosphate on T442 to stabilize the kinase active site conformation. To allow T442 trans-autophosphorylation, an entire thirty-three amino acid sequence from the kinase active site (indicated in yellow in Fig. 5C, Fig. 5D and in Movie Fig. 5CD), is proposed to exchange into another, already phosphorylated, PKR kinase domain in a face-to-face interaction ^28^. The interaction of ADAR1 dsRBDIII with the PKR kinase activation loop may prevent the activating phosphorylation of T442 by inhibiting the exchange of activation loops. This would prevent the spread of the phosphorylated, active state through clusters of PKR.

### ADAR1 binding to dsRNA and effects of ADAR1 dsRBDIII mutants on editing activity and PKR interaction

We wished to test the predicted ADAR1-PKR interaction using single residue changes in ADAR1 dsRBDIII. However, ADAR1 dsRBDIII is also expected to interact with the ADAR1 deaminase domain and to make contacts with dsRNA which play a critical role in accurate targeting of ADAR1 RNA editing sites. Therefore, we need to understand ADAR1 deaminase domain dimerization and dsRNA interactions also in order to interpret effects of dsRBDIII point mutants. In the ADAR2-dsRNA cocrystal structure ^29^, an asymmetric dimer of the deaminase domain is formed on dsRNA. The catalytic site and RNA interacting residues of the non-catalytic deaminase domain are blocked from interacting with dsRNA, as they are now part of the asymmetric deaminase domain dimerization interface. dsRBDII of the non-catalytic ADAR2 monomer (in hot pink), contacts dsRNA and positions its linked deaminase domain (in sandy brown) to hold the catalytic deaminase domain of the other monomer (also sandy brown), in position over the target adenosine (Fig. 5E, Movie Fig. 5E,F).

To examine how ADAR1 may use dsRBDIII in a similar way to bind to dsRNA we used AlphaFold Multimer to predict the asymmetric dimer structure of full length ADAR1 in the absence of dsRNA. We expect that the catalytic domains of the asymmetric deaminase domain dimers of ADARI and ADAR2 will bind very similarly to the dsRNA at the target adenosine so that they can flip the adenosine base out of the dsRNA into the deaminase active sites similarly. Therefore, we used the Chimera Matchmaker tool to superimpose the catalytic deaminase domain of the AF-predicted ADAR1 dimer (ADAR1 deaminase domains are in sky blue) onto the ADAR2 catalytic deaminase domain (ADAR2 deaminase domains in sandy brown) in the ADAR2-dsRNA cocrystal. The non-catalytic ADAR1 deaminase domain is then similarly positioned to that of ADAR2, showing that the dimerization of the deaminase domain occurs very similarly. The non-catalytic ADAR1 deaminase domain is tilted a little further away from the dsRNA than that of ADAR2 (Fig. S4B, Movie Fig. S4B). We then hid the ADAR2 protein and the N-terminal domains of ADAR1 to show the predicted ADAR1 deaminase domain plus dsRBDIII dimer and how it may interact with the dsRNA substrate (Fig. 5D, Movie Fig. 5C,D).

The predicted interaction of ADAR1 dsRBDIII (in orange) with the ADAR1 deaminase domain (Fig. 5D, Movie Fig. 5C,D), is very different from the ADAR2 dsRBDII interaction with the ADAR2 deaminase domain. ADAR2 dsRBDII (hot pink in Fig. 5E, Movie Fig. 5E,F), interacts with the ADAR2 deaminase domain in such a way as to line dsRBDIII up parallel to the dsRNA so that it can bind along the dsRNA in the canonical way that dsRBDs bind. ADAR1 dsRBDIII however is a protein-interacting, type B, dsRBD; it has an additional α1 helix, not present in ADAR2 dsRBDII ^30^. This enlarged alpha helical end of type B dsRBDs is often used for protein-protein interactions and in ADAR1 dsRBDIII this end of the dsRBD is predicted to interact with the ADAR1 deaminase domain (Fig. 5F, Movie Fig. 5E,F), particularly through the C-terminal part of helix α2 (Fig. S4C, Movie Fig. S4C). ADAR1 dsRBDIII now points across towards the dsRNA rather than along it. This AlphaFold model does not predict RNA interactions and it only gives a relative positioning of ADAR1 domains compared to dsRNA. However, of the three canonical dsRNA-contacting regions in dsRBDs^30^, only the second region at the tip of the beta sheet loop and the third SKKxAK region at the start of helix 3 in ADAR1 dsRBDIII come close to dsRNA (Fig. 5F, Movie Fig. 5E,F).

We identified predicted PKR-interacting residues in ADAR dsRBDIII (marked in magenta in Fig. 5C, Movie Fig. 5C,D and Fig. S4C, Movie Fig. S4C) that are away from the deaminase domain and should not affect dsRNA binding by impairing the ADAR1 dsRBDIII-deaminase domain interaction (Fig. S4C, Movie Fig. S4C). We introduced a single mutation, ADAR1 K777E, in a dsRBDIII residue usually involved in dsRNA-binding that is also predicted to be close to the PKR activation loop in the ADAR1-PKR complex (Fig. 5D, Movie Fig. 5C,D). ADAR1 K777E changes the first lysine in the third set of canonical dsRNA-contacting residues at the N-terminus of dsRBDIII helix α3 (SKKxAK), to a negatively charged glutamate. In canonical dsRNA-binding the SKKxAK residues usually bridge across the major groove, with the first lysine in the motif contacting phosphates on a different strand from lysines two and three. Surprisingly, when ADAR1 K777E is expressed from a transfected plasmid in HEK293 cells having a reporter in which luciferase is expressed when ADAR1 editing removes a stop codon, ADAR1 K777E reduces RNA editing activity only very slightly compared to wildtype ADAR (Fig. S5A). When assayed on *pri-miR 376a2* substrate coexpressed in HEK293 cells, editing at this site is also only slightly reduced compared to editing by ADAR1 p150 (Fig. S5B). This finding is not consistent with a canonical mode of dsRBDIII binding along dsRNA but it could be consistent with the ADAR1 dimer-dsRNA interaction model in Fig. 5D, Movie Fig. 5C, Movie Fig. 5CDD, if the interaction of the first lysine, K777, across the major groove that occurs in canonical binding along the dsRNA no longer occurs when ADARI dsRBDIII instead points towards one strand of the dsRNA. ADAR1 K777E expressed in A549 cells after plasmid transfection fails to interact with PKR in co-immunoprecipitation assays (Fig. 5B); suggesting that this residue may also participate in PKR contacts; although it is not predicted to contact PKR directly in the individual AF model shown, it is, nonetheless, close to PKR.

We also tested the ADAR1 R790E mutant, which changes an ADAR1 dsRBDIII residue which is more strongly predicted to contact PKR and which is located further along helix 3, away from RNA-contacting residues. The ADAR1 R790E mutant strongly reduces RNA editing activity on the luciferase substrate (Fig. S5A). When ADAR1 R790E activity is assayed on *pri-miR 376a2* substrate, editing at the +4 site is reduced even below background levels (Fig. S5B), suggesting that the ADAR1 R790E protein may still bind editing sites but the protein is altered and does not support editing. The ADAR1 R790E mutant also very strongly prevents ADAR1 co-immunoprecipitation with PKR (Fig. 5B). In the AF models of the dsRBDIII-PKR kinase domain interaction, lysine 790 reaches back along the alpha helix α3 and interacts with aspartate 786 in the same helix, suggesting that the R790E change removes a positive charge and adds a negative charge at this residue and may expose the aspartate to make a total charge change of minus three along this helix. The ADAR1 R790E mutant protein increased negative charge and alteration of protein structure might affect PKR interaction and also perhaps the helix α2 interaction with the deaminase domain. The ADAR1 3xEAA and ADAR1 dsRBDIII mutant effects are consistent with a requirement for ADAR1 dsRNA-binding for PKR interaction and are also consistent with our AF models for the ADAR1 PKR interaction and the ADAR1 dimer dsRNA-interaction.

## Discussion

These findings identify PKR as the second main innate immune sensor, after Mda5, that is aberrantly activated in *Adar Mavs* pups that are null mutants for Adar1 protein. *Adar Mavs* P14 pups show dramatic intestinal cell death, specifically in TACs in intestinal crypts, which virtually eliminates these cells. Normally, the TAC progenitor cells should differentiate and move up to maintain the absorptive intestinal villi by rapid replacement of villi cells as old villi cells die or are sloughed off. *Adar Mavs* pups are unable to maintain their intestinal villi and most are dead by P21. In *Adar Mavs EIF2αk2* triple mutant pups also lacking Pkr, early death of pups is substantially rescued, as is the death of intestinal crypt TACs and the loss of intestinal villar structure. Surprisingly, *Adar Mavs EIF2αk2* pups do show intestinal cell death again at P20 but cell death appears to be postponed beyond the TAC stage in the intestinal cell renewal process and is now distributed in basal regions of villi, allowing villar structure to be maintained.

Control of Pkr activity by Adar1 is mainly an editing-independent effect of a direct interaction between Adar1 dsRBDIII and the Pkr kinase domain on dsRNA. This ADAR1-PKR protein-protein interaction on dsRNA is a significant new mechanism of ADAR1 protein function. ADAR1 proteins define their target dsRNAs by binding to them and inhibiting PKR activation on them as well as by the editing of the dsRNA. A parallel report in bioArchiv also showed that removing Pkr rescues *Adar Ifih1* (Mda5) mutant mice ^31^.

Interaction of ADAR1 protein with PKR is likely to be part of a more extensive protein interaction network for control of innate immune sensor activation by proteins on dsRNA. PKR in particular is known to have many protein-protein interaction partners. ADAR1 dsRBDIII also engages in interactions with other innate immune sensor proteins, such as DICER ^32^. In *Drosophila*, Dicer2 also acts as a virus dsRNA-activated innate immune sensor ^33^ which leads to induction of antimicrobial peptide transcripts.

The success of AlphaFold in protein structure prediction is partly due to the limited number of different protein fold families ^34^; AlphaFold can also predict stable protein complexes and might also succeed in predicting transient protein interactions, particularly if there are recurring patterns in these. It has been suggested that protein domains, as well as having a limited number of protein folds, might also have an unexpectedly limited number of ways to interact with each other ^35^. For instance, we wondered if, since PKR dsRBDs interact with the PKR kinase domain to prevent PKR activation, an ADAR1 dsRBD might interact with the PKR kinase domain to exert a similar effect. The PKR dsRBDs and ADAR1 dsRBDIII are of different types and they use different parts of the domain for the interaction but both are positioned where they can interact with the dynamic activation loop of the PKR kinase domain. The AlphaFold-predicted interaction of ADAR1 dsRBDIII with the PKR kinase domain (Fig. 5D, Movie Fig. 5C, Movie Fig. 5CDD) places helix3 in a similar place on the kinase domain C-lobe to that of PKR dsRBDI helix1 in the AF-predicted human PKR monomer structure (Fig. 5C, Movie Fig. 5CD, Movie Fig. 5AB). We expect that dsRBDs of further proteins inhibiting PKR, such as DHX9 ^36^, will also interact with the PKR kinase domain.

Intestinal cell death has been previously reported in *Adar* mutants ^11,12,37^; however, we have examined this intestinal cell death in more detail. An *Adar* knockout specifically in *Lgr* intestinal stem cells shows a cell-autonomous role of Adar1 and generates the set of defects most similar to, and more severe than, those in our *Adar Mavs* mice ^21^. Adar1 may play a cancer-relevant, cell-autonomous role in maintaining the TAC cells, as *Adar* null mutants or tissue-specific *Adar* knockouts have also been shown to lead to loss of cancer stem cells^38^ and hematopoietic progenitor cells^3^. Proliferating stem cells require lin28 protein; ADAR1 prevents processing of *let-7* miRNA, which inhibits lin28 expression ^38^. In other *Adar* mutants that are not nulls, intestinal cell death seems delayed to later stages after TACs differentiate into villus cells ^12,37^.

Defects in *Adar Mavs* double mutant intestines are probably also partly due also to loss of Adar1 in EGCs that help maintain the stem cell niche in the intestinal crypt. Conditional *Adar* knockout in neural crest cells leads to loss of neural crest derivatives such as skin melanocytes, giving rise to unpigmented mice, Schwann cells myelinating peripheral neurons and probably also to loss of enteric glial cells ^39^. Some neural crest-derived enteric neurons and glia probably do survive in the *Adar* conditional knockout neural crest as these mice have normal villi up to their death at P5 ^39^. The *Adar Mavs* intestinal crypts also lack Paneth cells, which respond to Wnt signaling and help sustain crypt stem cells through additional Wnt signal production. Absence of Paneth cells or some other feedback from the aberrant TAC cells could cause the aberrant increase in Olfm4-positive true quiescent intestinal stem cells; TACs can also revert to being quiescent stem cells. Rapid loss of villi ^40^, similar to that seen in the *Adar Mavs* double mutant and accelerated apical migration of TAC cells and Paneth cells are also observed with specific knockout of E-Cadherin in mouse intestines. The Paneth cells are also abnormally large and may also have features of Goblet cells ^40^, like those we observe in P14 *Adar Mavs* intestines (Fig. 3A, C, D).

Our findings complement and extend several previous reports and also differ from them in some respects. Considering first those studies that used *Adar* null or *Adar^p^*^150^ mutant mice; defects in another *Adar Mavs* double mutant mouse also involve early death of pups and intestinal cell death ^12^. However, the cells that die are different from our *Adar Mavs* double null mutant; synchronous death of TAC progenitor cells was not observed and the intestinal cell death is milder and more like that seen in our *Adar Mavs EIF2αk2* mutant after twenty days. This other *Adar Mavs* double mutant is also rescued for pup survival and intestinal cell death by adding Zbp1 mutants removing Z RNA-binding protein 1^12^; ADAR1 also interacts with ZBP1 ^41^. Differences from our results may be mainly due to the newer *Adar* deletion mutant used in that report^12^, which is really a presumed *Adar^p^*^150^ null which retains expression of Adar1 p110. Mice with deletions in the *Adar* gene that remove or disrupt the deaminase domain while leaving N-terminal Adar1 coding sequences in place have been prone to generate unexpected truncated proteins with strong aberrant mutant phenotypes ^42,43^.

Mouse mutants in the Adar1 first Z-DNA/RNA binding domain, Zα have been studied as models for AGS. For example, Maurano et al ^11^ studied *Adar^P^*^195^*^A^*, which is equivalent to the AGS6 mutant *ADAR^P^*^193^*^A^* which generates full-length ADAR1 p150 protein defective in binding to Z RNA. These are excellent mouse models for AGS disease; however, available evidence presented indicates that the molecular mechanisms involved differ somewhat from those in *Adar* null mutants. The *Adar^Zα^* mutants give early death of mutant pups and intestinal defects only when heterozygous with *Adar* null or *Adar^p^*^150^–specific knockout mutants, suggesting the mutant phenotypes arise due to haploinsufficiency of the mutant protein. An *EIF2αk2* (Pkr) mutant rescues *Adar^P^*^195^*^A^* /*Adar^Δ^*^7–9^ and *Adar^P^*^195^*^A^* /*Adar^p^*^150^– mutant combinations by preventing aberrant induction of the integrated stress response^11^. We confirm the importance of PKR activation in *Adar Mavs* but do not observe a similarly strong aberrant activation of ISR transcripts in liver and other organs nor in *Adar Mavs* intestines; ISG expression in intestines is also only slightly activated in intestines. The stronger ISR induction in the *Adar^P^*^195^*^A^* /*Adar^p^*^150^– mice leading to defects similar to eIF2α mutant-mediated vanishing white matter syndrome, could reflect an even stronger aberrant activation of PKR than in our *Adar Mavs* double null mutants, perhaps due to some Adar1 mutant protein gain of function effect in *Adar^P^*^195^*^A^* /*Adar^Δ^*^7–9^ and *Adar^P^*^195^*^A^* /*Adar^p^*^150^–.

Several other reports show that *Adar1^Zα/-^* mutant combinations showing intestinal cell death and early death of pups, are rescued by adding a *Zbp1* mutation that postpones or reduces, but does not completely prevent, intestinal cell death ^12,37,41^. We observe that *Adar Mavs EIF2αk2* triple mutant intestines still show a delayed onset of cell death in intestinal villi at P20, so reducing and postponing cell death in intestines may allow maintenance of intestinal villi and allow pup survival. This may be the reason why either *EIF2αk2* or *Zbp1* mutations are able to rescue *Adar1^Zα/-^*mutant defects. Our findings also differ in detail from these other reports. Increased expression of Zbp1 was observed in *Adar^P^*^195^*^A^* combinations with other mutants and elsewhere ^12,41^; however, and we also do not observe very increased Zbp1 in *Adar Mavs* intestines. One report examined molecular mechanisms downstream of Zbp1 activation, i.e. effects on Ripk1/3 signaling, and found differences between *Adar* null mutants and *Adar1^Zα/-^* mutant combinations. *Adar* null mutants would simply eliminate all Adar1 interactions with other proteins but *Adar1^Zα/-^*mutant mice still express Adar1 p150 Zα mutant proteins which could exert some aberrant negative effect on an Adar1 protein interaction. ADAR1 Z RNA-binding is required to localize ADAR1 p150 to stress granules ^44^, close to where PKR also localizes. Thus, it would be informative if subcellular localizations of human ADAR1 P193E and Adar1 proteins in *Adar1^Zα/-^* mutant mouse cells would be examined.

The ADAR1-PKR interaction may be critical in the human brain, where *ADAR* mutants give rise to AGS symptoms and to striatal neurodegeneration. PKR protein has important roles in the brain, is regulated by many other proteins and can also be activated without dsRNA through other proteins such as PKRA and TARBP2. Aberrant activation of PKR in *Adar1^Zα/-^* mutant mice and in DYT16 dystonia patients, due to loss of the PKR regulatory protein PKRA, leads to massive death of neurons^45^.

We have demonstrated a novel mechanism whereby ADAR1 can inhibit PKR that is independent of RNA editing but requires dsRBDIII. This helps explain why mutational analyses of *Adar* in mice give complex results because the different domains of ADAR1 have evolved to regulate innate immunity; the Zα domain functions in binding to Z-RNA and possibly to ZBP1, the dsRBDs function in dsRNA-binding, the deaminase domain in RNA editing and now dsRBDIII makes an inhibitory interaction with PKR.

## Methods

### Mouse Breeding

Our *Adar^Δ^*^2–13^ null mutant mouse strain, which is also named *Adar^tm2Phs^* in the MGI database, lacks exons two to thirteen of *Adar*, encoding residues 11-1054, which comprise two Z RNA-binding domains, all three dsRNA-binding domains, and most of the catalytic domain of ADAR1, which are replaced with a *pgk-neo* gene. All of our current mouse strains have been generated after revival, using frozen sperm collected from our previous animals at Edinburgh University, to fertilize eggs from C57Bl6N females at the Czech Centre for Phenogenomics in Prague. Progeny mice were brought to the mouse facility at Masaryk University Brno, and bred together to produce strains with different mutant combinations in the C57Bl6N strain background. Our *Adar^Δ^*^2–13^, Mavs double mutant strain has been described previously ^1^.

The *EIF2αk2* mutant was generated in the Czech Centre for Phenogenomics at the Institute of Molecular Genetics ASCR using Cas9-mediated deletion of *EIF2αk2* exons 2 (containing the first AUG) to 5 coding for dsRNA-binding domains (amino acids: 1-165). Sequences of guide RNAs were Pi1B: 5’-GTGTTTCCAACCCACCAC<u>AGG</u> in the intron 1 and Pi5B: 5’-GGATCATTGTTGGTACAC<u>AGG</u> in the intron 5 (yielding 7,338-nt deletion). To produce guide RNAs, synthetic 128 nt guide RNA templates including T7 promoter, 18nt sgRNA and tracrRNA sequences were amplified with T7 and tracrRNA primers. Guide RNAs were produced *in vitro* with the Ambion mMESSAGE mMACHINE T7 Transcription Kit (Thermo Fisher, #AM1344), and purified with mirPremier™ microRNA Isolation Kit Sigma-Aldrich, #SNC50). The Cas9 mRNA was synthesized from pSpCas9-puro plasmid with Ambion mMESSAGE mMACHINE T7 Transcription Kit, and purified with the RNeasy Mini kit (Qiagen, #74104). A sample for microinjection was prepared by mixing two guide RNAs in water (25 ng/μl for each) together with Cas9 mRNA (100 ng/μl). Five pl of the mixture were microinjected into male pronuclei of C57Bl/6 zygotes and transferred into pseudo-pregnant recipient mice. PCR genotyping was performed on tail biopsies from four-weeks-old animals. A positive founder which transmitted the mutant allele to F_1_ was back-crossed with C57Bl/6NCrl animals for at least five generations before using in experiments.

The knock-out allele was detected with mPKR_i1_Fwd: 5′-GCCTTGTTTTGACCATAAATGCCG and mPKR_E6_Rev: 5′-GTGACAACGCTAGAGGATGTTCCG primers giving a 552 bp product (wild-type allele is too long to be amplified). Wild-type allele was detected with mPKR_i1_Fwd and mPKR_E2_Rev: 5′-GTAGAAACCTGGGGTATCACTGGC primers yielding a 361 bp product.

### Collection of mouse embryos

Five C57BL/6 females in fertile age and with the required genotype were group-housed in the same cage for at least 10 days to synchronize their estrous cycle. Females were then exposed to pheromone secreted in male urine by introduction of soiled bedding from male’s cage for 3 nights to increase the chances of synchronized estrous cycle. Only mouse females that will be in proestrus or estrus were chosen for mating and one to one breeding was performed during late evening. The following morning the vaginal plug was checked and midday on the day when the mating plug was detected is counted as embryonic day E0.5. Embryos were then collected, genotyped by taking their tail and head and body were snap frozen for protein or RNA extraction or fixed in 4% PFA for histology.

### Immunoblotting

Mouse tissues were lysed in lysis buffer (100 mM KCl, 5 mM MgCl_2_, 10% Glycerol, Tris-HCl 20mM [pH 8], Tween 20 0.1%) supplemented with protease inhibitors (Halt™ Protease Inhibitor Cocktail, EDTA-Free (100X) #78439), PMSF and phosphatase inhibitor (Halt™ Phosphatase Inhibitor Cocktail (100X) #78427) using ceramic beads in a tissue homogenizer grinding machine at 5,000rpm for 30s for 2 cycles. HEK 293 T-Rex cells were lysed in lysis buffer (0.5% NP40, 50mM HEPES, 5mM EDTA, 150mM NaCl, 1X Protease and 1X Phosphatase Inhibitors) and centrifuged at 12,000g for 20 minutes. Lysates were then clarified centrifuging them for 20 minutes at 12,500xg at 4°C. The protein concentration was measured by BCA (Pierce™ BCA Protein Assay Kit #23225). 40 μg (intestinal tissue) of protein lysate was separated by SDS-PAGE, transferred on a nitrocellulose membrane and blotted at 95V for 1.15h at 4°C. The membranes were blocked with 5% BSA or 5% dry milk in TBS-T for 1h at room temperature and incubated with a primary antibody for an overnight at the following dilution: Adar1 1:200 (to detect mouse Adar, Santa Cruz, #sc-73408), Pkr 1:2,000 (Abcam #ab184257 [EPR19374]), p-eIF2α (Ser51) 1:1,000 (Cell Signaling, #3398), eIF2α 1:1,000 (Cell signaling, #9722), Rig-I 1:500 (Cell Signaling, [D14G6] #3743), Mda5 1:1000 (Cell Signaling [D74E4],#5321), Gapdh 1:20,000 (Proteintech, #60004-1-Ig). Adar1 1:3000 (to detect human ADAR, Antibodies-online.com #ABIN2855100), HA tag Rabbit 1:1000 (Cell signaling #3724), HA tag mouse 1:2000 (antibodies.com #A85278) were also used. A secondary antibody was used in TBS-T with a dilution of 1:5,000 for Anti-Mouse (Thermo Fisher, Goat anti-Mouse IgG, #31430) and 1:80,000 for Anti-Rabbit antibody (Merck Anti-Rabbit IgG, #A0545). Clarity Western ECL Substrate (Bio-Rad, #1705061) was used as a substrate for detection of horseradish peroxidase (HRP) enzyme conjugates and a UVITEC ChemiDoc imaging was used to expose the membrane as well as Odyssey® M (Li-COR) machine.

### Immunofluorescence

The proximal intestine was flushed with cold PBS and fixed in 4% PFA for 24h at 4°C. After 3 washes of 10 minutes with cold PBS at room temperature, the tissue was embedded in OCT overnight at –80°C. The tissue was then cryosectioned with a thickness of 10 μm at –22°C, and sections air-dried onto the slides at room temperature to maximize their adherence. The slides were stored at –80 °C. Before performing the staining, the slides were air-dried for 30 minutes and rehydrated by washing them 3 times for 10 minutes with the washing buffer (PBS + 0.1% Triton X100). To minimize non-specific binding of primary and secondary antibodies, blocking buffer (PBS + 0.1% Triton X100 + 1% BSA + 0.15% glycine) was used for 1h and 30 minutes at room temperature. Sections were incubated in a humidified chamber overnight with a primary antibody at the following dilution: E-cadherin 1:200 (Invitrogen #XA3425032), GFAP 1:200 (Agilent #ZO334), Lysozyme 1:150 (Agilent, EC 3.2.1.17 #A0099), SMA 1:300 (Abcam, #ab5694), Anti-beta III Tubulin 1:400 (Abcam, #ab52623), CD45 1:200 (Thermo Fisher, #AB_467251). The slides were then rinsed 4 times with the washing buffer for 15 minutes and incubated with the secondary antibody at 1:500 dilution in blocking solution for 1h at room temperature (Rabbit 555, Invitrogen, #AB_162543; Rat Cy3, Jackson, #AB_2340667). Slides were rinsed with washing buffer 4 times for 15 minutes and then washed 5 minutes with PBS. Mounting was performed with liquid mounting media with DAPI (VECTASHIELD® PLUS Antifade Mounting Medium with DAPI, Vector Laboratories #H-1200-10).

Olfm4 staining was performed on wax embedded sections. After a 3 paraffin removal steps using xylene, sequential ethanol incubations (100%, 95%, 80%, 60%) follow to rehydrate the tissues. Antigen retrieval step was performed using a pressure cooker (110 degrees, 15 minutes) and a citrate buffer (pH 6.0). The slides were cooled to reach RT and a 25 minutes peroxidase blocking followed (0.3% H_2_O_2_ in methanol) to prevent false positive signal. After washing the slides 4 times in TBS (pH 7.4) a 1h-blocking step (5% goat serum, 1% BSA, 0.05% Tween, 1xTBS pH 7.4) was performed to prevent non-specific binding. Sections were incubated overnight at 4°C in a moist chamber (Olfm4, Proteintech 1:100). After washing the slides with TBS + 0.01% Triton for 5 minutes, they were rinsed 2 times in TBS. Incubation with a Secondary antibody Anti-rabbit Alexa 488 GAR-488 A11034 Invitrogen (1:500, 1hour at RT) in 5% goat serum, 1% BSA in TBS followed. After washing the slides in TBS, ProLong Gold Antifade Mountant with DAPI was used and a cover glass placed.

### Tissue preparation

Intestines were cut from a P14 pups. Stomach and caecum were discarded and the small intestine was flushed first with cold PBS and then with cold Bouin’s fixative (50% ethanol, 5% acetic acid in dH_2_O). The intestine was then divided into 3 equal parts (proximal, medial and distal), open longitudinally while kept in Bouin’s fixative and rolled following the Swiss Roll technique ^46^. Formalin fixation was performed in 4% PFA for 24h. Tissues were then washed with cold PBS 3 times for 10 minutes and stored in 70% EtOH. The tissue was dehydrated in increased percentages of EtOH solutions: 70% EtOH for 45 minutes, 80% EtOH for 60 minutes, 96% EtOH for 60 minutes, 2 times 100% ethanol for 60 minutes. Later ethanol was exchanged with xylene in the following sequence: xylene for 45 minutes, xylene for 60 minutes, xylene for 75 minutes. Finally, xylene was exchanged with paraffin in a vacuum oven set for 58 °C: paraffin for 60 minutes, paraffin for 75 minutes, paraffin for 90 minutes. The tissue was embedded in fresh new paraffin in the desired orientation and left to solidify. 2.5 μm thick paraffin sections were cut using a microtome, collected onto a positively charged slide and dried overnight at 37 °C. After cooling to room temperature, the slides were used for staining.

## Immunohistochemistry

### Ki67 staining

Tissue sections were deparaffinized by submerging them 3 times in xylene for 5 minutes. Xylene was removed by placing the tissues in 100% EtOH two times for 5 minutes. Sections were rehydrated by sequential incubation in 95%, 80% and 60% EtOH, 5 minutes each. Slides were rinsed in distilled water for 5 minutes and then an antigen retrieval step was performed by placing the slides in EDTA buffer, pH 9 (DCS Innovative Diagnostik-Systeme #EL010C500), in a water bath at 98°C for 30 minutes. Slides were left to cool in the buffer at room temperature for 30 minutes, rinsed twice in distilled water for 5 minutes and washed in PBS for 5 minutes. To minimize the non-specific binding, a blocking step with 5% Goat serum, 0.1% Tween in PBS was performed for 1h at room temperature in a humidified chamber. Slides were incubated overnight with anti-Ki-67 1:100 (Zytomed systems, #RBK027) antibody at 4°C in a humidified chamber, then washed 3 times in PBS for 5 minutes. Endogenous peroxidase was blocked by submerging the slides in freshly prepared 3% H_2_O_2_ in PBS for 15 minutes at room temperature. Three five-minute washes with PBS were performed and the slides were then incubated for 30 minutes at room temperature with the SignalStain® Boost IHC Detection Reagent (HRP, Rabbit #8114, Cell Signaling) and, after one washing in PBS, they were subsequently incubated with DAB working solution (SignalStain® DAB Substrate, #8059, Cell Signaling) at room temperature until staining developed (1-10 minutes). Slides were washed in distilled water for 5 minutes. Sections were dehydrated by sequential five-minute incubations with 60%, 80%, 95% and 100% ethanol. After three five-minute washes in xylene, sections were mounted with Pertex, covered with a coverglass and air-dried under the hood.

### H&E staining

Tissue sections were deparaffinized in 3 xylene washes of 5 minutes followed by rehydration steps of 5 minutes each in: 100% EtOH, 80% EtOH, 70% EtOH. Slides were washed in dH_2_O for 5 minutes and stained with Hematoxylin for 10 minutes. After a 10 minute wash in warm water, the slides were stained in Eosin for 5 minutes and washed in dH_2_O for 5 minutes. Sections were dehydrated by sequential incubation with 70%, 80%, 100% EtOH, 5 minutes each. After 3 washes in Xylene of 5 minutes each, the slides were mounted with Pertex, covered with a cover glass and air-dried under the hood.

### TUNEL Assay

To detect cell death in intestinal tissues a terminal deoxynucleotidyl transferase dUTP mediated nick-end labeling (TUNEL) assay was performed following the manufacturer’s protocol (Manufacturer). The intestinal tissue was fixed in 4% PFA and paraffin-embedded. 3 µm sections were cut and labeled with a TUNEL reaction mixture containing terminal deoxynucleotidyl transferase and nucleotides including tetramethylrhodamine-labeled dUTP. When the protocol was concluded the slides were checked by fluorescence microscopy. DAPI staining appears as blue dots marking the nuclei whereas TUNEL staining appears as green dots indicating the presence of degraded DNA.

### Sequencing of microbiome

Quality of DNA was determined by gel electrophoresis and concentration was assesed spectrophotometrically with a microplate reader (Synergy Mx, BioTek, USA). For identification of bacteria presented in the samples, the sequencing of 16S rRNA gene was performed. Extracted DNA was used as a template in amplicon PCR to target the hypervariable region V4 of the bacterial 16S rRNA. The library was prepared according to the instructions for Illumina 16S Metagenomic sequencing Library Preparation protocol with some deviations described below. Each PCR was performed with the primer pair consisting of Illumina overhang nucleotide sequences, an inner tag and gene-specific sequences ^47^. The Illumina overhang served to ligate the Illumina index and adapter. Each inner tag, i.e. a unique sequence of 7–9 bp, was designed to differentiate samples into groups. The total reaction volume of PCR was 30 µl consisting of 15 μl Q5 HighFidelity 2x MM (BioLabs, New England), 1.5 μl of each 10 μM primer, 9 μl of PCR water and 3 μl of template. The cycling parameters included initial denaturation at 98°C for 30 s, followed by 30 cycles of 10 s denaturation at 98°C, 15 s annealing at 55°C and 30s extension at 72°C, followed by final extension at 72°C for 2 min. The amplified PCR products were determined by gel electrophoresis. PCR clean-up was performed with SPRIselect beads (Beckman Coulter Genomics). Samples with different inner tags were equimolarly pooled based on fluorometrically measured concentration using Qubit® dsDNA HS Assay Kit (Invitrogen™, USA) and microplate reader (Synergy Mx, BioTek, USA). Pools were used as a template for a second PCR with Nextera XT indexes (Illumina, USA). Differently indexed samples were equimolarly pooled based on fluorometrically measured concentration as before. The prepared library was checked with the three methods: qPCR using LightCycler 480 Instrument (Roche, USA) and KAPA Library Quantification Complete Kit (Roche, USA); 2100 Bioanalyzer Instrument with the D1000 ScreenTape kit (Agilent Technologies, USA) and microplate reader with dsDNA HS Assay Kit. Concentration was measured shortly prior sequencing. The final library was diluted to a concentration of 8 pM and 20 % of PhiX DNA (Illumina, USA) was added. Sequencing was performed with the Miseq reagent kit V2 with a MiSeq instrument according to the manufacturer’s instructions (Illumina, USA).

### RNA extraction and cDNA reverse transcription

Total RNA was obtained from mouse tissues using RNeasy Micro Kit (Qiagen #74004). Sample integrity and quantity was checked with the use of a TapeStation (RNA Screen Tape, Agilent). 5 µg of total RNA was reverse transcribed into cDNA with RevertAid Reverse Transcriptase kit, (Thermo Fisher, #EP0442), using random hexamers.

### Quantitative RT-PCR

RT-PCR was performed with FastStart Universal Syber Green Master (Rox) (Roche, #FSUSGMMRO). 45 cycles of amplification were performed 95°C for 10 sec and 60°C or 58°C for 20 seconds, 72°C for 30 seconds using LightCycler 480 II System (Roche). Gene-specific primers and 50ng of cDNA were used for the amplification. The comparative CT Method (ΔΔ*C*_T_) was used for genes relative quantification. Results were normalized to the mRNA expression of Hprt1. The primers used are in Table 2.

### Site-directed PCR mutagenesis of selected amino acids

Primers containing the mutations were designed and purchased from Generi Biotech and are listed in Table 2. The method used previously described ^48^. A 20µl PCR reaction with 10ng of pDONR221 plasmid containing ADAR1p150 as template, was performed with Phusion™ High-Fidelity DNA Polymerase (2U/µL) (Thermo Fisher, #F-530XL). The PCR conditions were set according to manufacturer’s instructions and the number of cycles was decreased to 19. 10µl of the PCR product was electrophoresed on a 0.8% agarose gel to confirm the amplification of the desired product. The remaining 10µl of the PCR reaction was digested with 0.5µl of *Dpn*1 enzyme 10U/µL (Thermo Fisher, #ER1701) at 37°C for 1 hour, followed by inactivation of the enzyme at 80°C for 20 minutes. 2µl from each sample was used to transform 100µl of TOP10 competent *E.coli* cells. The mixture was incubated on ice for 30 minutes, heat shocked at 42°C for 90 seconds, followed with 5 minutes on ice and then mixed with 1ml of LB medium and incubated at 37°C for 1 hour. The cells were then centrifuged at 5,000 rpm for 2 mins, pellet resuspended and 150µl was plated on agar plate with ampicillin resistance. Plates were left in the incubator at 37°C overnight. Colonies were then picked and plasmids were isolated with the Exprep™ Plasmid SV midi kit (GeneAll, #101-201) as per manufacturer’s instructions. Samples were then Sanger-sequenced by Eurofins Scientific to confirm the presence of the desired mutation. Gateway LR reaction with the Gateway™ LR Clonase™ II Enzyme mix (Thermo Fisher, #11791020) was performed to transfer ADAR1p150 containing the desired mutation from pDONR221 plasmid to the destination vector, p-II-Strep-HA-N tag ^49^, according to manufacturer’s instructions. The entire LR reaction was used to transform TOP10 competent cells, bacterial colonies were picked and plasmids were isolated with the Midi kit. Successful transfer of mutated ADAR1p150 into the destination vector was confirmed by Sanger-sequencing.

### Immunoprecipitation

A549 cells were transfected with constructs (GFP, ADAR1p150 wt, ADAR1p150 EAA, ADAR1p150 777, ADAR1p150 790) as per the Lipofectamine 3000 (Thermo Fisher, # L3000015) protocol. 4 hours later, IFNα (2000IU/ml) was added. After 24 hours, cells were lysed with lysis buffer (150mM NaCl, 50mM Tris-HCl pH 8, 1mM EDTA pH 8, 10% Glycerol, 0.5% NP-40) vortexed every 5 mins for 30 mins, and centrifuged at 18,000 × *g* at 4°C for 20 min. The supernatant was quantified and 1mg of protein was precleared for 1 hour with 20µl of Dynabeads^TM^ Protein G (Thermo Fisher, #10003D) for each sample. Samples were then incubated with 3µl each of PKR (abcam, [EPR19374] #ab184257) and IgG (Cell Signalling Technology, #2729) antibodies, overnight at 4°C. The next day, proteins were immunoprecipitated with 30µl of beads per sample for 1 hour at 4°C. Flow-through was collected and protein-bound beads were rigorously washed with wash buffer (200mM NaCl, 50mM Tris-HCl pH 8, 1mM EDTA pH 8, 10% Glycerol, 0.5% NP-40) every 10 mins for an hour. Beads were boiled at 95°C for 7 mins in Laemmli Buffer and were subjected to immunoblotting against their respective antibodies on nitrocellulose membrane. Clarity Western ECL Substrate was used as a substrate for detection of horseradish peroxidase (HRP) enzyme conjugates and a UVITEC ChemiDoc imaging was used for visualization.

### A-to-I editing activity assay

A-to-I editing activity was measured as previously described ^4^ as well as using a dual luciferase reporter sequence (adapted from ^50^ commercially synthesized by GenScript within 5’ attB1 and 3’ attB2 gateway cloning sites in pUC57 plasmid. The A-to-I editing reporter sequence contains a single open reading frame encoding mammalian optimized Firefly luciferase containing a N-terminal PEST destabilization domain (luc2P), followed by a 72 nucleotide fragment of *GluA2* sequence containing a known editing site within an in-frame stop codon (U<u>A</u>G) which when edited is converted to a tryptophan codon (U<u>G</u>G), followed by mammalian optimized nanoluciferase containing an N-terminal PEST destabilization domain (NlucP). ADAR activity results in read-through at the A-to-I edited stop codon and expression of nanoluciferase. Therefore, A-to-I editing levels are determined by the ratio of nanoluciferase to Firefly luciferase signal. For positive control, the in-frame stop codon (UAG) was been converted to tryptophan (UGG), giving expected signal for 100% editing by ADAR. For negative control, 18 codons of the *GluA2* sequence are removed from the region complementary to the UAG codon, preventing formation of dsRNA structure necessary for editing by ADAR, giving expected signal for 0% editing by ADAR. The A-to-I editing dual luciferase reporter was inserted into pCDNA3.2 V5-DEST plasmid (Thermo Fisher Cat. #1248019) by Gateway cloning (Thermo Fisher) following manufacturer’s instructions.

HEK293 cells were co-transfected with 1:1 ratio of cDNA3.1 expressing wither ADAR1p150 or mutations in ADAR1p150 and A-to-I editing dual luciferase reporter, 48 hours following transfection, signal was detected using Nano-Glo Dual-Luciferase Reporter Assay System (Promega Cat. #1610) following manufacturer’s instructions, and luminescence was measured on a Spark 10M multimode plate reader (Tecan, Mannedorf, Switzerland). The relative level of A-to-I RNA editing was calculated by ADAR transfected (A-to-I editing reporter [Nanoluciferase/Firefly Luciferase] – Negative control reporter [Nanoluciferase/Firefly Luciferase]) ^51^ / ADAR endogenous (A-to-I editing reporter [Nanoluciferase/Firefly Luciferase] – Negative control reporter [Nanoluciferase/Firefly Luciferase]).

### dsRBD-PKR kinase domain interaction predictions using AlphaFold and AlphaFold Multimer

The human PKR predicted monomer structure was taken as a downloaded PDB file from the AlphaFold Structure Database ^25,34^ and annotated and colored using PyMol. The predictions of the Adar1 dsRBDIII-PKR kinase domain interaction were made using an AlphaFold Multimer ^26^ installation on the Czech Supercomputer Server in Brno with default settings and generation of five or ten top models for each interaction. The sequences used for input can be seen by Display Sequence on the PyMol or Chimera Session files provided with the Supplementary Materials. PyMol was used to annotate the models and colour different regions and Chimera was used for the Matchmaker tool to superimpose interaction models for dsRBDIII-PKR interactions and ADAR1 and ADAR2 dimers. Models are presented as TIFs for each panel in Figure 5, as .mov movie files in the Supplementary material and also as portable session files in the Supplementary Materials. The ADAR2-dsRNA complex image is made from PDB file pdb8e0f using Chimera

### Software used for data analysis

GraphPad Prism was used to realize the Kaplan Meier curve and t-test analyses. ImageJ was used for the densiometric analysis of the immunoblots.

## Statistical analyses

The acquired images were analyzed with FIJI software. Cells that tested positive for TUNEL assay and were labeled with Alexa 488 were separated from the background by applying the Moments algorithm ^52^. Then, the number of particles that had a TUNEL positive foci above a threshold (T) was counted in a square area measuring 250 microns in length using the Sampling window plugin. Statistical significance is marked as *p < 0.05, **p < 0.01, ***p < 0.001, ****p < 0.0001 respectively

The area of the intestinal crypt base, which was indicated by the Olfm4 positive signal, was separated from the background with MaxEnthropy threshold algorithm, and the Olfm4 signal area was measured. For analysis, more than 50 areas were used. The statistical analysis, consisting of a one-way ANOVA followed by Kruskal-Wallis one-way analysis of variance, was performed using GraphPad Prism (version 9). Statistical significance is marked as *p < 0.05, **p < 0.01, ***p < 0.001, ****p < 0.0001 respectively

2-ΔΔCT algorithm was used to analyze the relative changes in gene expression after performing the RT PCR. These values were graphed after being normalized to the housekeeping gene Hprt1 and to one reference Wt value. Statistical analysis performed with one-way ANOVA followed by Kruskal-Wallis one-way analysis of variance, performed with GraphPad Prism software (version 9). Statistical significance is marked as *p < 0.05, **p < 0.01, ***p < 0.001, ****p < 0.0001 respectively. A minimum of 3 and a maximum of 5 different biological replicas and 3 technical replicas per each gene were used.

Microbiome Samples from this project were sequenced on the Illumina MiSeq NGS sequencing platform and the resulting reads were processed by the following preprocessing pipeline. The first step was demultiplexing of reads in sequencing pools into individual samples. This was conducted by an in-house tool written in Python 3. Each sample was identified by a predefined sequence of nucleotides (tag) located at the beginning of each read. This sequence was 7-9bp long and had to be located within first 30 nucleotides of a given read. The demultiplexing process was conducted at the same time for both reads in each pair. If any of the reads in pair had unexpected tag sequence, or no tag sequence at all, the whole pair was discarded. After demultiplexing, the tag sequences and further 30 nucleotides were removed from the beginning of each read. This was done in order to ensure, that no adapter sequences would be present in subsequent analysis.

The next step in the pipeline was trimming of low-quality end of each read. All reads in trimming have had fixed number of N last nucleotides removed. Because, forward and reverse reads tend to have a very different quality profiles, the trimming length N was always determined for forward and reverse reads separately. The determination of the trimming length was conducted in each run according to the same rule. Let X denote the first position in read, at which, an average first quartile of the phred score in subsequent 5 sequence positions was below 20 and let L denote the expected read length after demultiplexing, then, the trimming length N was determined as N = L – X. For example, if the expected length of reads in a given run was 206 bp after demultiplexing, and the first point where the average first quartile of phred score of 5 subsequent sequence positions was 191, the resulting trimming length was 206 – 191 = 15 bp. As a result, last 15 bp were trimmed from each read.

Following the quality trimming, pairs, where any read contained the nucleotide N, were discarded. Pairs, in which any of the reads was shorter than a predetermined threshold, were also discarded. Given the expected read length L and the trimming length N, this threshold was always calculated as T = L – N – 15. In the case of the previous example, the length threshold would be set to 206 – 15 – 15 = 176 bp. Like in the case of the trimming length, the length threshold was calculated for forward and reverse reads separately. If any of the reads in pair was shorter than its respective length filtering threshold, the whole pair was discarded. Both, N filtering and length filtering, were performed by an in-house tool written in Python 3.

After the reads were trimmed, the possible contaminants were inspected. Reads were mapped onto the human reference genome (GRCh37) and the phix174 reference genome. This was performed with the bowtie 2 NGS aligner ^53^. Reads, which mapped onto any of these genomes were discarded from further analysis.

Next, forward and reverse reads were denoised using the DADA2 amplicon denoinsing R package ^54^. This was done in order to cope with the sequencing and PCR-derived error. Following denoising, the forward and reverse reads were joined using the fastq-join read joining utility ^55^. In order to be joined, reads in pair had to have an overlap of at least 20bp with no mismatches allowed. Pairs in which this was not the case were discarded. As a final step, chimeric sequences were removed from the joined reads using the remove Bimera function of the DADA2 R package.

## Animal Studies

The ethical approval to perform the mouse studies is in license MSMT-32428/2016-3 and MSMT-28797/2021-4 from the Czech Ministry of Education, Youth and Sports.

## Supporting information

Fig. S4A ADAR1 dsRBDIII and PKR 10 predictions

Fig. S4B ADAR1 fl dimer Matchmaker to ADAR2 on dsRNA

Fig. S4C ADAR1 dsRBDIIII helix2 and linker contacts to first helix of deaminase domain

## Acknowledgements

We are very thankful to Katarína Marečková and Aleš Hampl for assistance with processing mouse sections for imaging. We are very grateful to Petr Svoboda for the mutant *EIF2αk2* mice. We thank Karel Kubicek for help with AlphaFold and with the local AlphaFold Multimer installation. The Imaging Core Facilities of CEITEC Masaryk University is gratefully acknowledged as well as ARCC for use of a slide scanner. DV was supported by GAČR 20-11101S and KS by GAČR 19-16963S. This work was also supported by a grant from the Czech Science Foundation GAČR 21-27329X to MAO’C, by Agence Nationale de la Recherche RNAediting-inPNS (ANR-21-CE12-0020) research grants to NB, work of R.M. was supported by RVO 68378050-KAV-NPUI, and by Czech Academy of Sciences RVO68378050 and LM2018126 for the Czech Centre for Phenogenomics provided by MEYS CR to RS. Authors thankthe RECETOX Research Infrastructure (No LM2018121) financed by the Ministry of Education, Youth and Sports, and the Operational Programme Research, Development, and Education (the CETOCOEN EXCELLENCE project No. CZ.02.1.01/0.0/0.0/17_043/0009632) for microbiome sequencing and analysis.

## Authors contributions

Ketty Sinigaglia: tissue preparation, immunohistochemistry, immunofluorescence, immunoblotting, RT-PCR, preparation of figures.

Anna Cherian: Immunoblotting and immunoprecipitation

Dragana Vukic: Immunoblotting

Janka Melicherova: animal husbandry

Pavla Linhartova: animal husbandry

Qiupei Du: RNA editing assays

Lisa Zerad: immunohistochemistry, immunofluorescence

Stanislav Stejskal: immunofluorescence

Radek Malik: generation of *EIF2αk2* mutant

Jan Prochazka: generation of *EIF2αk2* mutant

Nadège Bondurand: supervision, immunohistochemistry, immunofluorescence

Radislav Sedlacek: revival of *Adar Mavs* strains

Mary A. O’Connell: supervision and writing manuscript

Liam P. Keegan: designed experiments, supervision and writing manuscript

