## Supplementary Information for "Aberrant activation of the innate immune sensor PKR by self dsRNA is prevented by direct interaction with ADAR1"

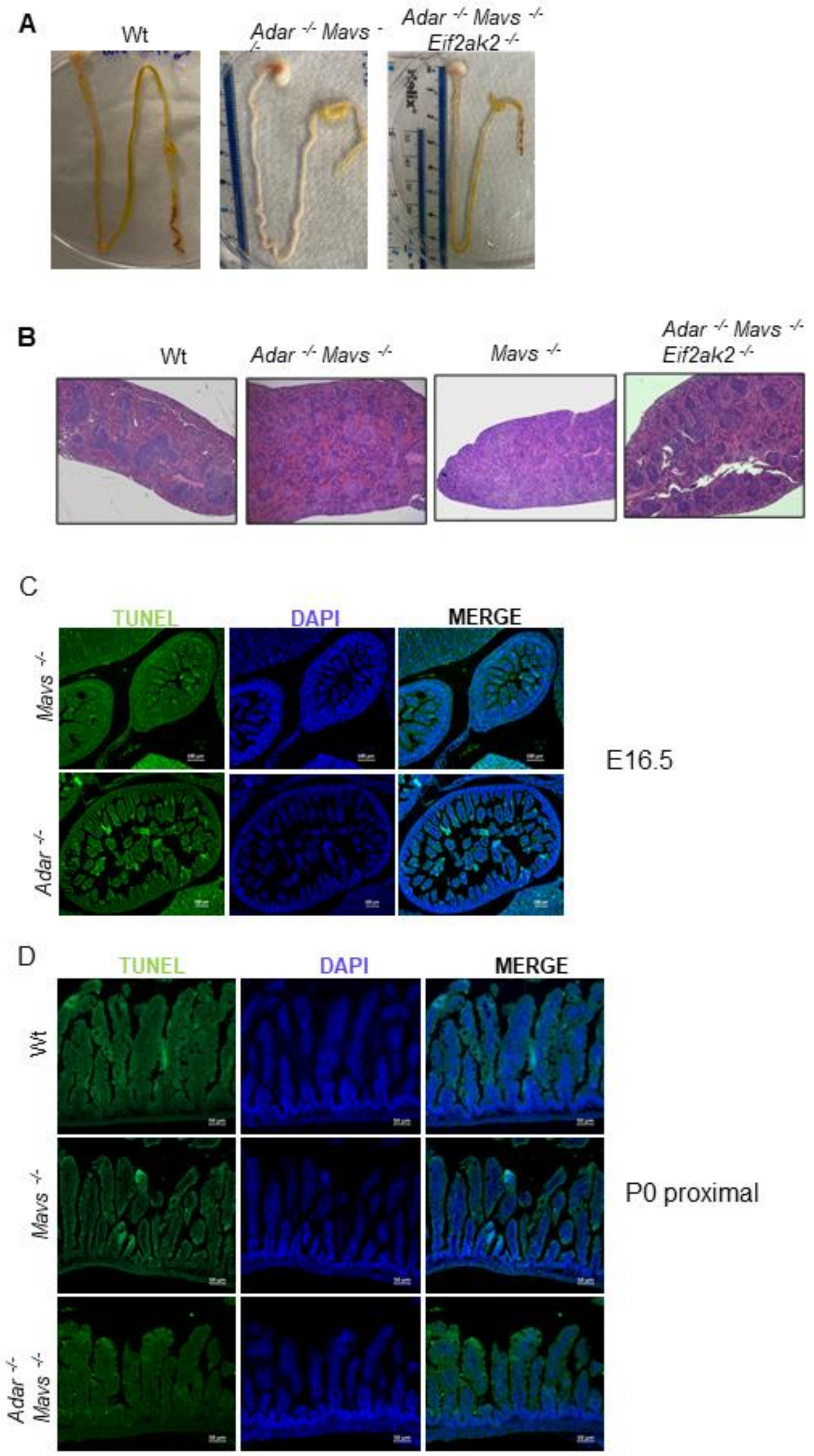


**Supplementary Figure 1. Intestinal defects arise postnatally. A.** *Adar Mavs* intestine length appears similar to *Adar Mavs Eif2ak2* length and the ratio between body size and intestine length is normal. **B.** Hematoxylin and eosin staining shows the spleen morphology with loss of white matter (which stains red with eosin) in the *Adar Mavs* double mutant pup and the different morphologies in two *Adar Mavs Eif2ak2* pups at P14. Diffuse spleen congestion and simultaneous white pulp hypoplasia are evident in the double mutant and are still present in one of the 2 rescued pups. **C.** TUNEL-positive cell death is absent embryonically at E16.5. Scale bar is 100 µm. **D.** TUNEL-positive cell death is absent in proximal intestine at birth, P0. TUNEL-positive cells are shown in green and nuclei are stained with DAPI (blue).  Scale bar is 50 µm.

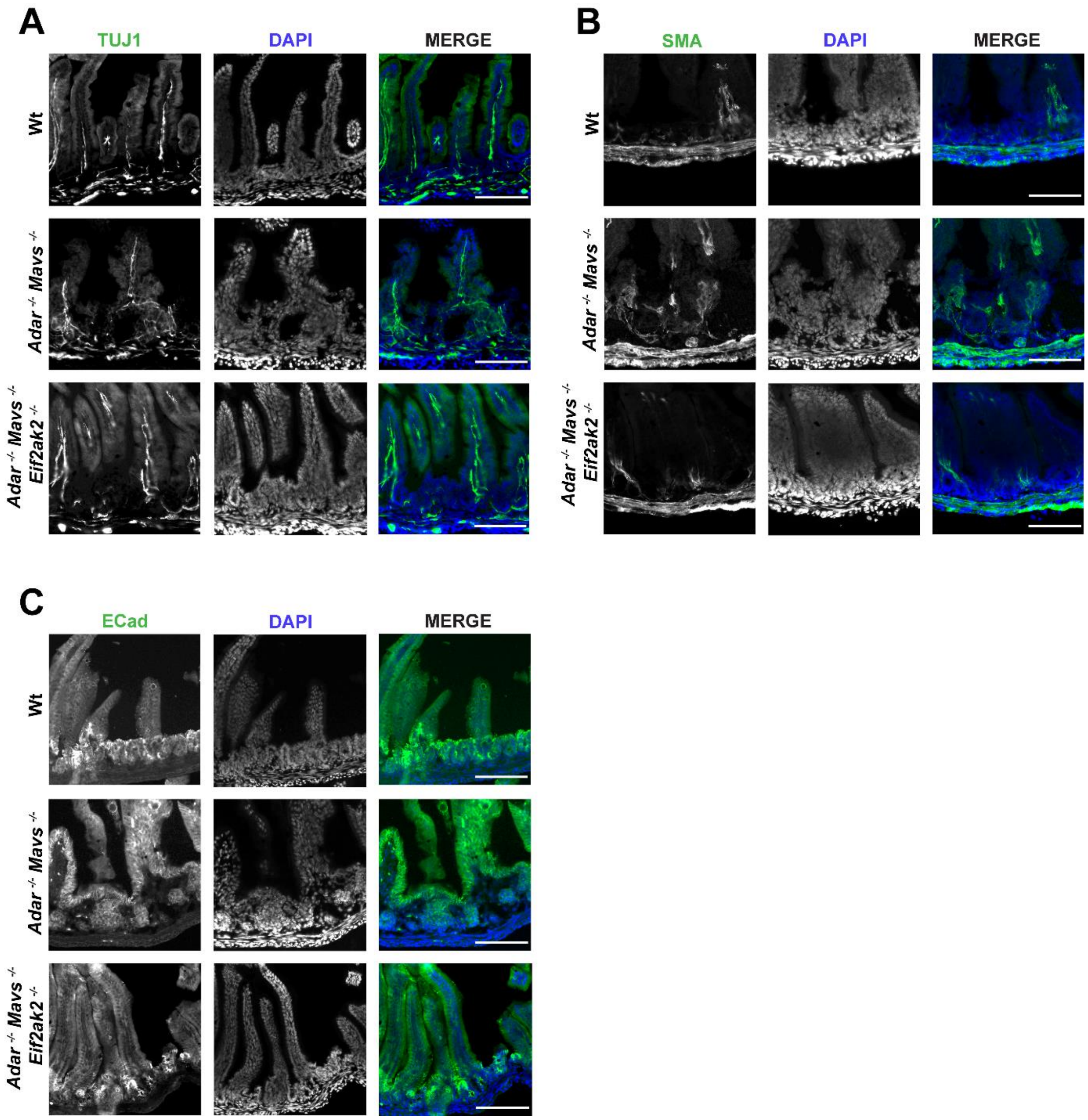


**Supplementary Figure 2. Normal enteric neurons and muscle cells in *Adar Mavs* intestines and rescue of *Adar Mavs* aberrant intestinal E-cadherin staining but not spleen defects the *Adar Mavs Eif2ak2* mice. A.** Neuron-specific B-tubulin (TUJ1) appears less organized in *Adar Mavs* P14 proximal small intestine due to disrupted villi morphology. **B.** Smooth muscle (SMA) is not damaged in *Adar Mavs* P14 proximal intestine. **C.** Anti-E-cadherin staining, DAPI staining and merged images show organized expression in all epithelial cells of the villi in wild-type and in *Adar Mavs Eif2ak2* intestines*;* anti-E-cadherin staining is disorganized in basal regions of the villi in *Adar Mavs*)**.** Two biological replicas were used in the panels above. Scale bar is 100 μm in all panels.

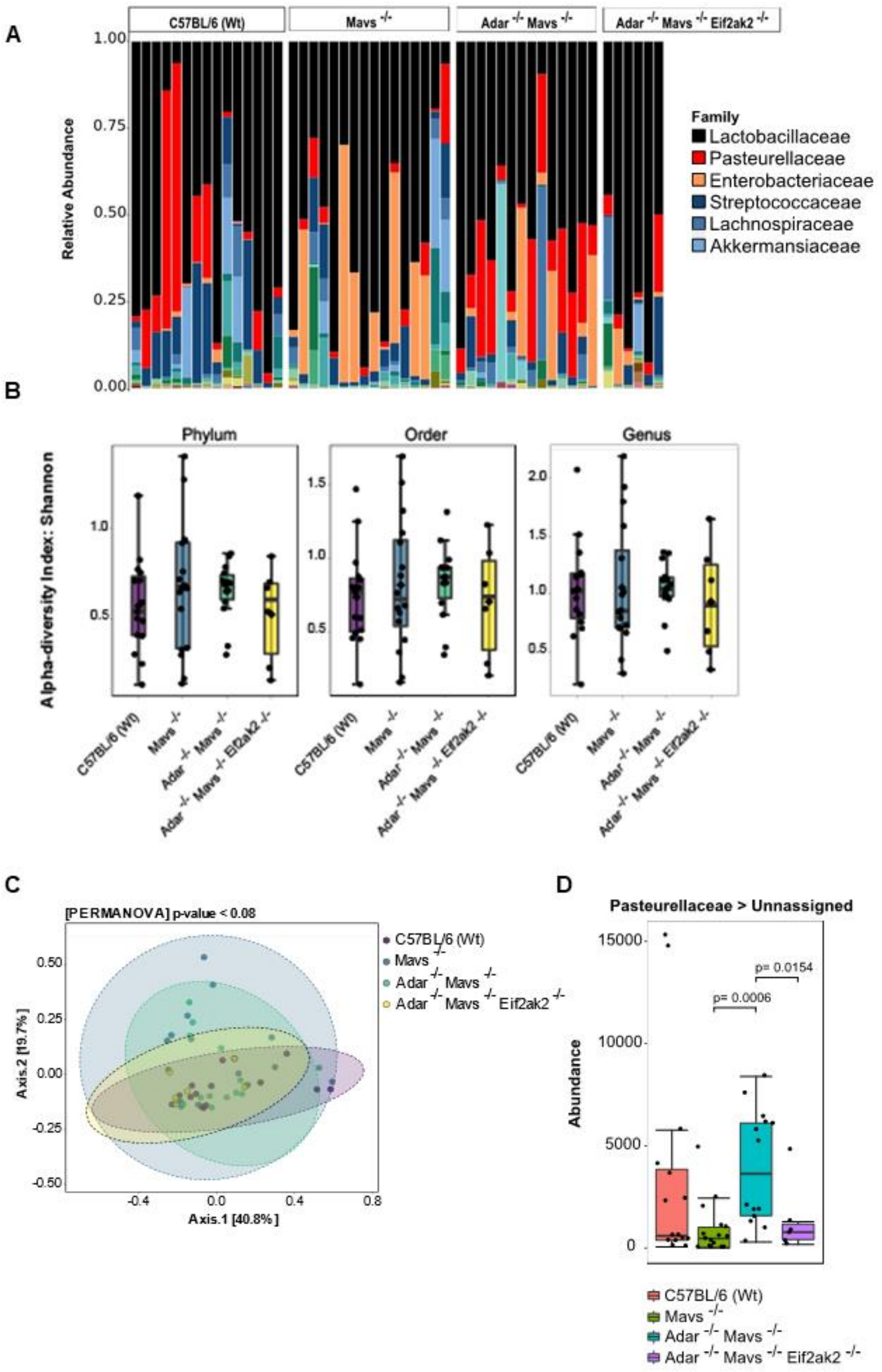


**Supplementary Figure 3. Sequencing of microbiome. A.** Abundance plots of 16S rRNA amplicon sequencing results. Relative abundance of different bacteria taxa at the Genus level in the mouse pup small intestine (duodenum, jejunum, ileum) at P14. Legend shows the most abundant taxa. Sample size: Wt = 15, *Mavs*$^{-/-}$ = 16; *Adar*$^{-/-}$*Mavs*$^{-/-}$ = 14; *Adar*$^{-/-}$*Mavs*$^{-/-}$*Eif2ak2*$^{-/-}$ = 6. **B.** Comparison of gut microbiota Alpha diversity between each group at Phylum, Order and Genus level. Alpha diversity indicates the diversity of microorganisms within a single sample. It quantifies how many different types of microorganism are present and how evenly they are distributed. Shannon Diversity Index was used to measure Alpha diversity. Analysis of Variance (ANOVA) was performed to statistically analyze the differences between the means. **C.** Differences in gut community Beta diversity indicate the differences in species composition. Bray-Curtis Dissimilarity method was used to calculate dissimilarities between the groups considered. It provides a value between 0 and 1 where 0 indicates identical communities and 1 indicates dissimilar communities. Statistical method used Permutational multivariate analysis of variance (PERMANOVA) was used as statistical method. **D.** *Pasteurellaceae* abundance comparison between the different mouse genotypes.

**Supplementary Figure 4. - movies**

**Chimera Matchmaker superpositions of AlphaFold Multimer Model predictions and PyMol image of ADAR1 dsRBDIII residues interacting with the deaminase domain. A.** Chimera Matchmaker superpositions of ADAR1 dsRBDIII-PKR kinase domain AlphaFold Multimer protein interaction models. Each model shows the protein chains in a different color. The PKR kinase chains of the models have been superposed onto that in model0. The positions of the dsRBDIII then differ mainly by rotation around the N-terminus of dsRBDIII helix3 and the beta sheet loop which are in similar positions on PKR in three of the five models. See also Movie Fig. S4A. **B.** Chimera Matchmaker superposition of the catalytic deaminase domain of the AlphaFold Multimer-predicted dimer of full length ADAR1 onto the catalytic deaminase domain of ADAR2 bound to dsRNA. The N-terminal domains of ADAR1 have been hidden and only the top predicted model for the ADAR1 dimer is used here to avoid too much crowding. The dsRBDs bind dsRNA very differently in ADAR2 and ADAR1. See also Movie Fig. S4B. **C.** Pymol-annotated portion of the ADAR1 full-length dimer prediction showing dsRBDIII (in orange) and the linker after it (in white) contacting the deaminase domain (sky blue). The mutated dsRBDIII residues that may interact with PKR kinase domain are shown in magenta and these are far for the deaminase domain contacts. See also Movie Fig. S4C.

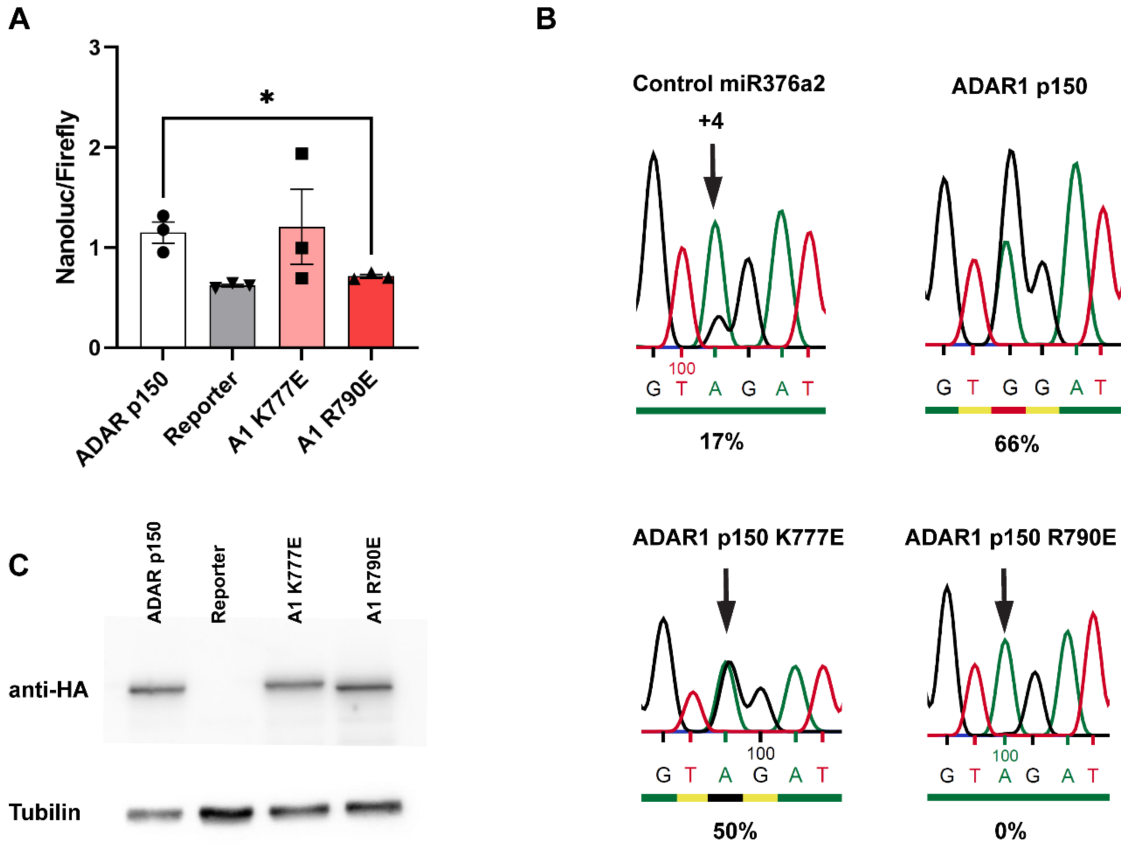


**Supplementary Figure 5. Effects of ADAR1 K777E and ADAR1 R790E mutations on ADAR1 RNA editing and dsRNA binding. A. Above.** Levels of luciferase expression in HEK293 cotransfected with an editing reporter cells and ADAR1 mutant or wildtype protein are expressed from transfected plasmids. The HEK293 transiently transfected RNA editing reporter expresses an RNA transcript having a dsRNA hairpin with an editable stop codon preceding the luciferase ORF. There is fifty percent editing of the reporter alone in the absence of any additional ADAR due to the background expression of ADARs in the HEK293 cells. Below. Immunoblot showing levels of wildtype and mutant ADAR1 proteins expressed in transfected HEK293 cells. **B.** Sequence chromatograms showing levels of editing at site +4 in the *pri-miR 376-a2* substrate transcript when expressed from a plasmid co-transfected into HEK293 cells with plasmids expressing wildtype or mutant ADAR1 proteins; adenosine is in green and guanosine is in black. Percentage editing is given and the background without additional ADAR is 17%. ADAR1 K777E edits almost as well as wildtype ADAR1 and

ADAR1 R790E suppresses the background editing, indicating that it does not edit but does bind to dsRNA.

**Supplementary Table 1. Primers used for qPCR and mutagenesis**

| qPCR primers | Forward Primer | Reverse Primer |
|---|---|---|
| m-OasL1 (60°C) | CCAACAATGTGGCAGAAGGC | GTGCTCTCTTCACCCTCCAG |
| m-Oas3 (60°C) | AGACCCTAGCTGAGGACCTG | CAGAAGACCCACCCTTGACC |
| m-Oas1-g (60°C) | ATGGAGCACGGACTCAGGAGCAT | TCACAGCAGGATACATGTCCAGTTC |
| m-Cxcl10 (58°C) | AAGTGCTGCCGTCATTTTCTGCCTC | CTTGATGGTCTTAGATTCCGG |
| m-Ifit3 (60°C) | TTTCCCATCAGCACAGAAAC | TTCAGCTGTGGAAGGATCG |
| m-Isg15 (60°C) | GGTGTCCGTGACTAACTCCAT | TGGAAAGGGTAAGACCGTCCT |
| m-Mx1 (58°C) | GACCATAGGGGTCTTGACCAA | AGACTTGCTCTTTCTGAAAAGCC |
| m-Atf4 (58°C) | CCTGAACAGCGAAGTGTTGG | TGGAGAACCCATGAGGTTTCAA |
| m-Cdkn1a (58°C) | TCGCTGTCTTGCACTCTGGTGT | CCAATCTGCGCTTGGAGTGATAG |
| m-Hprt1 (58°C/60°C) | TGGATACAGGCCAGACTTTGTT | CAGATTCAACTTGCGCTCATC |
| Primers for mutagenesis | | |
| ADAR1p150 K777E | AGCAAG<u>GAA</u>CAAGGCAAGCAGGA AGCAGCAGAT | GCCTTG<u>TTC</u>CTTGCTGTGTGCGCA GACGGC |
| ADAR1 p150 R790E | GCTCTC<u>GAA</u>GTCTTGATTGGGGAG AACGAGAAG | CAAGAC<u>TTC</u>GAGAGCCGCATCT GCTGCTTC |
